# Diverse gut pathogens exploit the host engulfment pathway via a conserved mechanism

**DOI:** 10.1101/2023.04.09.536168

**Authors:** Mahitha Shree Anandachar, Suchismita Roy, Saptarshi Sinha, Agyekum Boadi, Stella-Rita Ibeawuchi, Hobie Gementera, Alicia Amamoto, Gajanan D. Katkar, Pradipta Ghosh

**Author notes:** **CORRESPONDING AUTHORS: Gajanan D. Katkar, Ph.D.;** Departments of Cellular and Molecular Medicine, University of California San Diego, CA 92093. : **Email:**, **Pradipta Ghosh, M.D.;** Professor, Departments of Medicine and Cellular and Molecular Medicine, University of California San Diego, CA 92093. : **Email:**.

## Abstract

Macrophages clear infections by engulfing and digesting pathogens within phagolysosomes. Pathogens escape this fate by engaging in a molecular arms race; they use *WxxxE* motif-containing “effector” proteins to subvert the host cells they invade and seek refuge within protective vacuoles. Here we define the host component of the molecular arms race as an evolutionarily conserved polar ‘hotspot’ on the PH-domain of ELMO1 (*E*ngu*l*fment and Cell *Mo*tility1), which is targeted by diverse *WxxxE*-effectors. Using homology modeling and site-directed mutagenesis, we show that a lysine triad within the ‘patch’ directly binds all *WxxxE*-effectors tested: SifA (*Salmonella)*, IpgB1 and IpgB2 (*Shigella*), and Map (enteropathogenic *E. coli*). Using an integrated SifA•host protein-protein interaction (PPI) network, *in-silico* network perturbation, and functional studies we show that the major consequences of preventing SifA•ELMO1 interaction are reduced Rac1 activity and microbial invasion. That multiple effectors of diverse structure, function, and sequence bind the same hotpot on ELMO1 suggests that the *WxxxE*-effector(s)•ELMO1 interface is a convergence point of intrusion detection and/or host vulnerability. We conclude that the interface may represent the fault line in co-evolved molecular adaptations between pathogens and the host and its disruption may serve as a therapeutic strategy.

**GRAPHICAL ABSTRACT:** *In brief:* This work defines the nature of a conserved molecular interface, assembled between diverse W*xxxE* motif-containing effector proteins encoded by gut pathogens and the host innate immune sensor, ELMO1, via which pathogens exploit the host’s engulfment machinery. 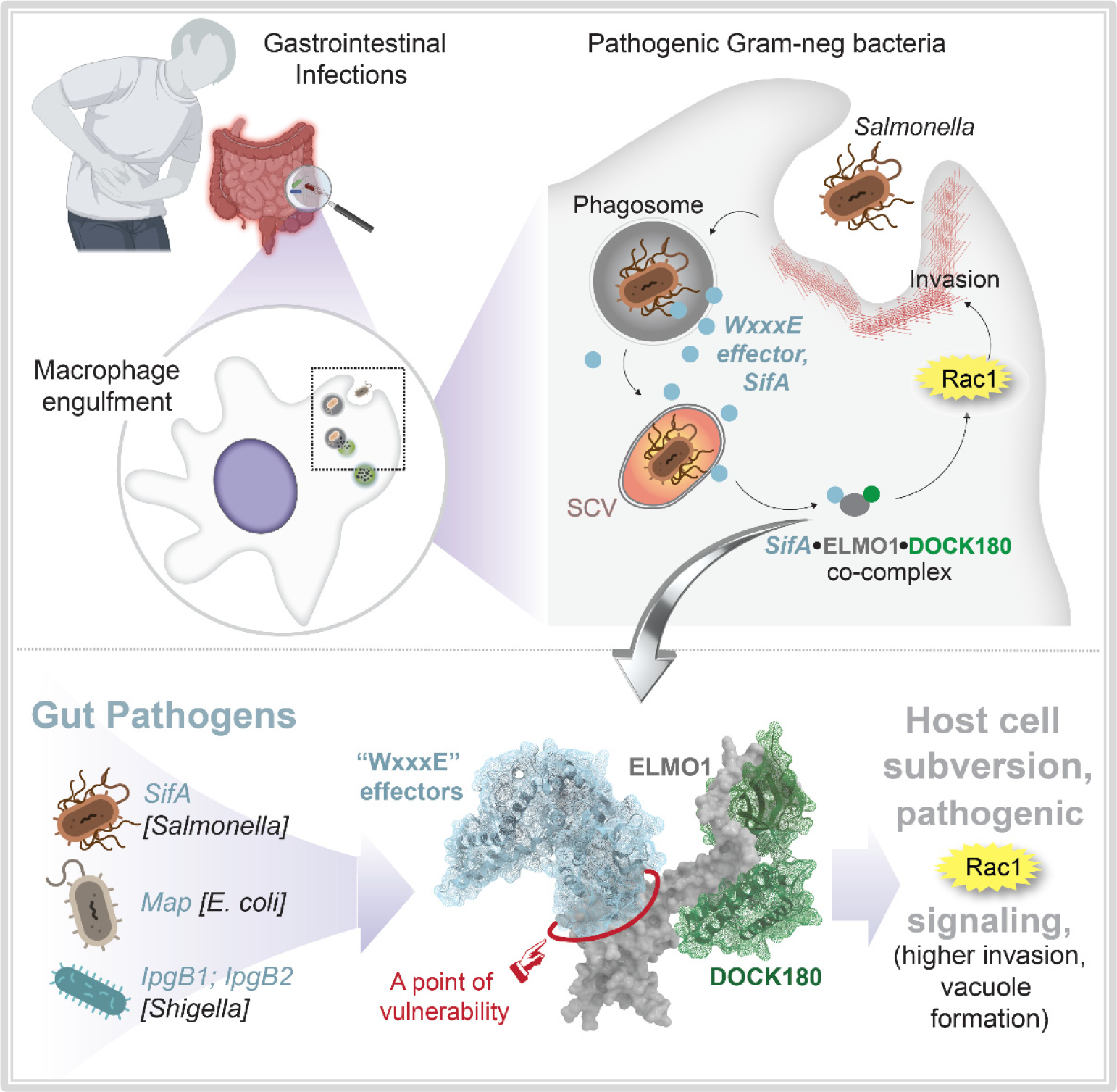

## INTRODUCTION

Enteric pathogens such as *Salmonella* rely upon their virulence factors to invade and replicate within host cells. Upon invasion, they seek refuge within a modified phagosome-like structure, the *Salmonella*-containing vacuole (SCV) (1), within which they survive, even replicate, or simply persist in a dormant-like state (2). Both invasion and SCV formation require the delivery of microbial effector proteins via Type III Secretion Systems (T3SSs) into the host cell; they both require the cooperation of a subverted host cell whose phagolysosomal signaling and membrane trafficking pathways are manipulated to mount very dynamic and extensive membrane remodeling and actin rearrangement (1). Thus, three key aspects facilitating *Salmonella* pathogenesis are (i) vacuole formation for refuge, (ii) delivery of effector proteins via T3SSs to interfere and/or manipulate the host system, facilitating (iii) more bacterial invasion. These mechanisms of pathogenesis are shared also among other enteric pathogens such as enteropathogenic *E. coli* EHECs, EPECs, and *Shigella. E. coli* rely upon T3SS effectors--e.g., Map, EspH, and EspF--to form *E. coli*-containing vacuoles (ECVs) (3) and manipulate the host cell form pedestals (4), filopodia, or microspikes for invasion, whereas *Shigella* rely upon T3SS effectors, e.g., IpgB1/2, to form vacuoles (5) and manipulate host cells into forming membrane ruffles for orchestrating what is known as “the trigger mechanism of entry” (6-9). Similar mechanisms are also used by *Yersinia* (10) and *Campylobacter* (11) to subvert host epithelial cells.

Regardless of the diversity of the pathogens, their equally diverse T3SS injectosomes, or the repertoire of effectors (reviewed in (12)), the host actin cytoskeleton has emerged as the dynamic hub in a microbe-induced circuitry of Ras-superfamily GTPases [Ras homolog family member A (RhoA), Ras-related C3 botulinum toxin substrate 1 (Rac1) and Cell division control (Cdc) protein-42] (13). Diverse microbes converge upon and exploit the host circuitry to mount pathogenic signaling, escape lysosomal clearance by seeking refuge in vacuoles, invade host cells and alter inflammatory response. As for mechanism(s) for such convergence, the ability of a *WxxxE* motif-containing family of effectors to directly activate host GTPases was reported first (14). By activating host Rho GTPases, the WxxxE effectors subvert actin dynamics (15). SopE, IpgB1 and EspT trigger membrane ruffles (16-18), IpgB2 and EspM trigger stress fibers (19) and Map trigger filopodia (20) via activation of Rac1, RhoA and Cdc42 respectively. *E*ngu*l*fment and cell *Mo*tility protein 1 (ELMO1) was identified subsequently as a *WxxxE* effector-interacting host protein (21). Three independent groups, each using ELMO1-knockout animals that were infected with different pathogens (*Shigella (6), Salmonella* (21)and *E. coli* (22)), have implicated the *WxxxE*•ELMO1 interaction in the augmentation of the actin•GTPase circuitry via the well-established ELMO1•DOCK180 (Dedicator of cytokinesis)→Rac1 axis (23-25). Within this signaling cascade, ELMO1•Dock180 is a bipartite guanine nucleotide exchange factor (GEF) for the monomeric GTPase Rac1 (26), but is also capable of activating Cdc42 (27) and RhoA (28).

Despite these insights, key questions remained unanswered, e.g., how do the *WxxxE*-effectors, which are unique to gut pathogens (21) (yet to be found in commensals (29)) converge on one host macro-molecular complex (the ELMO1•Dock180→RhoGTPases) to subvert host actin dynamics. Because the *WxxxE*-effectors are structurally and functionally diverse except for the *WxxxE*-motif, which is their defining and unifying feature, initial studies hypothesized that this motif could be the mechanism of such convergence; but four structural studies revealed otherwise (26, 30-32). These studies of SifA, IpgB, and Map structures and a EspM2 model showed that Trp (W) and Glu (E) within the *WxxxE-*motif are ‘structural residues’ that are positioned around the junction of the two 3-α-helix bundles and maintain the conformation of a ‘catalytic loop’ through hydrophobic contacts with surrounding residues. Consistent with these conclusions, conserved substitutions (W→Y and E→D; in EpsM (19)) did not alter stability or functions, whereas W→A or E→A substitutions make the protein highly unstable (33) and render it non-functional (34). With the *WxxxE-*motif ‘ruled out’ as the potential contact site for convergence, the basis for how diverse *WxxxE*-effectors may bind ELMO1 and induce convergent pathogenic signaling via the ELMO1•DOCK axis remains unknown. Using a transdisciplinary and multi-scale approach that spans structural models as well as protein-protein interaction networks, here we reveal a surprisingly conserved molecular mechanism for how diverse pathogens use their *WxxxE*-effectors to hijack the ELMO1•DOCK•Rac1 axis via a singular point of vulnerability on ELMO1.

## RESULTS AND DISCUSSION

### ELMO1 and SifA cooperate during vacuole formation

Prior work has separately implicated both ELMO1(35) and SifA (32, 36, 37) in *Salmonella* pathogenesis. While SifA has been directly implicated in the formation of SCVs (32, 37), ELMO1 was shown to impact bacterial colonization, dissemination, and inflammatory cytokines *in vivo* (21). We asked if both proteins are required for SCV formation. We stably depleted ELMO1 in J774 macrophages by short hairpin (sh)RNA (>99% depletion compared to controls; **Figure 1A**), infected them with either the WT (SL) or a mutant *Salmonella* strain that lacks *SifA* (*ΔSifA*), and then assessed the ultrastructure of the SCVs by transmission electron microscopy (TEM) (see workflow; **Figure 1B**). Bacteria were observed as either intact within vacuoles, free in cytosol, or partially digested within fused lytic compartments, as reported previously (38) (**Figure 1C-D**). When assessed for the completeness of the vacuolar wall (see basis for quantification; **Figure 1E**), WT *Salmonella* formed complete SCVs at a significantly lower rate in ELMO-depleted macrophages compared to controls (13% vs 35% **Figure 1F, H**). Absence of SifA impaired SCV biogenesis regardless of the presence or absence of ELMO1 (**Figure 1G, H**). Findings demonstrate that both ELMO1 and SifA are required for SCV formation and suggest cooperativity between the two proteins during SCV biogenesis.

**Figure 1:**
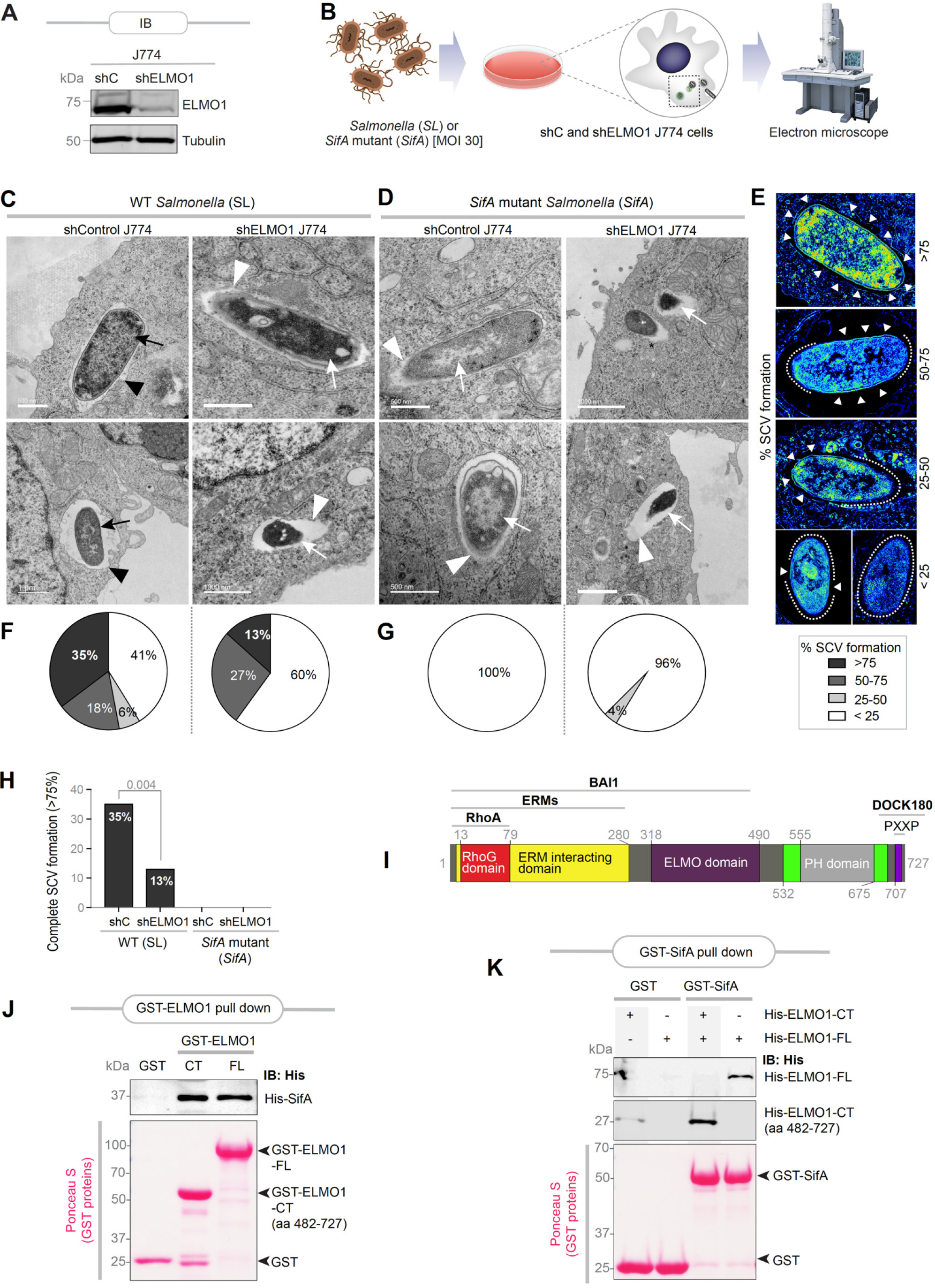
ELMO1 and SifA interact directly and cooperate to promote SCV formation. **A**: Immunoblot of control (shC) and ELMO1-depleted (shELMO1) J774 murine macrophages. **B**: Schematic shows workflow for TEM on J774 macrophages infected with either *Salmonella* (SL) or a *SifA*-deleted variant strain (SifA) of the same. **C-D**: Representative electron micrographs of control (shC) or ELMO1-depleted (shELMO1) J774 macrophages, infected with *S. typhimurium* (SL or *SifA*; 30 moi), at 6 h post-infection. Examples of bacteria within vacuoles (black arrowheads) and cytosolic bacteria (white arrowheads) are indicated. Bacteria that are either Intact (black arrows) or partially degraded upon fusion with lytic compartments (white arrows) are also indicated. Scale bars are placed at the lower left corner of each micrograph. **E-G:** A montage (E) of pseudo-colored micrographs of SCVs at various stages of formation is shown. The presence (arrowheads) or absence (interrupted lines) of detectable vacuolar membrane are marked. Pie charts display the percentage of bacteria at each stage of SCV formation (F-G) encountered in C-D. **H:** Bar graph display the percentage of complete (representing >75% in E) SCV formation in C-D. **I**: A domain map of ELMO1 with major interacting partners. ERM proteins (ezrin, radixin and moesin); PH, pleckstrin homology; BAI1, Brain-specific angiogenesis inhibitor 1 (*ADGRB1*). **J:** Recombinant His-SifA (∼5 μg) was used in a pulldown assay with immobilized bacterially expressed recombinant GST-tagged full-length (FL) or a C-terminal domain (CT; aa 482-727) of ELMO1 or GST alone (control). Bound SifA was visualized by immunoblot using anti-His (SifA) antibody. GST proteins are visualized by Ponceau S staining. **K**: Recombinant His-tagged CT (aa 482-727) or full-length (FL) ELMO1 proteins (∼3 μg) were used in a pulldown assay with immobilized bacterially expressed recombinant full-length GST-SifA or GST alone (control). Bound ELMO1 proteins were visualized by immunoblot using anti-His (SifA) antibody. GST proteins are visualized by Ponceau S staining.

### SifA directly binds the C-terminus of ELMO1

We next asked if SifA binds ELMO1; the latter is a multi-modular protein with several known interacting partners (summarized in **Figure 1I**). We compared head-to-head equimolar amounts of GST-tagged full length vs. a C-terminal fragment (aa 482-727) of ELMO1 [which contains a pleckstrin-like homology domain (PHD)-; **Figure 1I**] for their ability of to bind a His-tagged recombinant SifA protein. The C-terminal PH-domain was prioritized because of two reasons: (i) A recent domain mapping effort using fragments of ELMO1 on cell lysates expressing SifA had ruled out contributions of the N-terminal domain in mediating this interaction (21); and (ii) SifA directly binds another PH-domain, SKIP, forming 1:1 complex at micromolar dissociation constant, the structural basis for which has been resolved (26, 39). We found that both full length ELMO1 and its C-terminal fragment can bind His-SifA (**Figure 1J**). Binding was also observed when the bait and prey proteins were swapped, such that immobilized GST-SifA was tested for its ability to bind His-ELMO1 proteins (**Figure 1K**). Because interactions occurred between recombinant proteins purified to >95% purity, we conclude that the SifA•ELMO1 interaction is direct. Because SifA bound both the full length and the C-terminal fragment of ELMO1 to a similar extent, we conclude that the C-terminus of ELMO1 is sufficient for the interaction.

### SifA binds to an evolutionarily conserved lysine hotspot on ELMO1’s PH domain

To gain insights into the nature of the SifA•ELMO1 interface, we leveraged two previously resolved structures of a SifA•SKIP-PHD co-complex (32) and ELMO1-PHD to build a homology model of SifA•ELMO1-PHD complex (see **Fig.S1A-B** for workflow and *Methods*). The resultant model helped draw three key important conclusions: (i) the resolved structure of SifA•SKIP (32) and the model for SifA•ELMO1 were very similar, and hence, the specific recognition of SifA by both SKIP-PHD and ELMO1-PHD was predicted to be mediated through a large network of contacts (**Figure 2A**), primarily electrostatic in nature (**Fig.S1C-D**); (ii) the tryptophan (W197, deeply buried within the hydrophobic core of SifA) and glutamate (E201) within the *WxxxE*-motif (red residues; **Fig. S2**), which are essential for protein stability, but dispensable for binding SKIP (26) are likely to be nonessential also for ELMO1; and (iii) the amino acids deemed essential for the assembly of the SifA•ELMO1 interface were: a pair of hydrophobic residues, Leucine (L)130 and Methionine (M)131 on SifA and a triad of polar lysine residues within the β5-β6 loop of ELMO1 (K620, K626 and K628) (**Figure 2A-B**). An alignment of the sequences of ELMO1-PHD and SKIP-PHD showed the lysine triad in ELMO1 corresponds to the corresponding contact sites on SKIP for SifA in the resolved complex(26) (**Figure 2C**; **Fig. S3A**). A full catalog of both inter- and intramolecular contact sites (see **Supplemental Information 1**) revealed how each lysine within the lysine triad in the β5-β6 loop of ELMO1 contributes uniquely to generate the electrostatic attractions that stabilize the SifA•ELMO1 complex (**Figure 2B**): (i) K628 primarily establishes intermolecular electrostatic contacts with L130 and M131 on SifA; (ii) K626 mediates intermolecular interaction by engaging K132 and also via charge-neutralizing salt-bridges with Asp(D)117 on SifA. It also mediates intramolecular interactions with D621 within the β5-β6 loop of ELMO1; (iii) K620 primarily engages in intramolecular contact with two other residues within the β5-β6 loop, L631 and L638, thus stabilizing the loop. Thus, all 3 lysines within the triad appeared important: While K628 and K626 establish strong electrostatic interactions with SifA, K626 and K620 stabilize the β5-β6 loop that contains the lysine triad.

**Figure 2:**
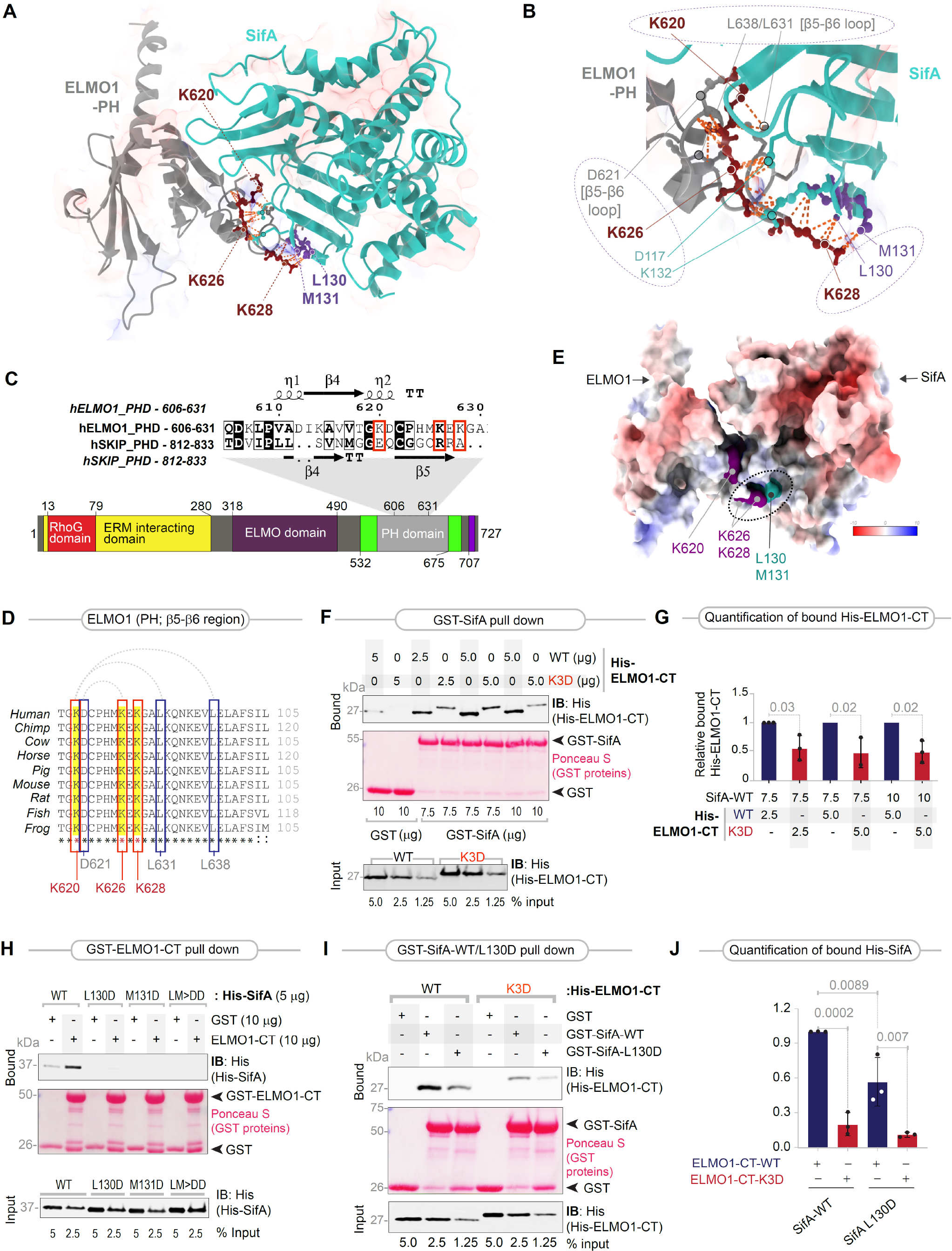
Characterization of the ELMO1•SifA interface. **A**. Homology model of the ELMO1(gray)•SifA(turquoise) complex, generated by superimposing the solved structure of the ELMO1-PH (PDB:2VSZ) on that of the SKIP-PH in complex with SifA (PDB:3CXB). The executive root mean square deviation (RMSD) of the model is 1.528 (see **Fig. S1** for the workflow used to generate the homology models and select the fittest model with the most optimal parameters). The model is annotated with key amino acids that are predicted to form the interface. On ELMO1, these amino acids were identified as an evolutionarily conserved polar hotspot of three lysines within the β5-β6 loop, and their key intra- and intermolecular contacts on SifA are annotated. See **Supplemental Information 1** for a complete catalog of the inter- and intramolecular contacts of the highlighted residues. The distance between the residues calculated for contacts were within 4.0 A. See **Fig. S2** for the position of the WxxxE motif relative to the ELMO1•SifA interface. **B**. A magnified view of the key residues participating at the ELMO1 (gray)•SifA (turquoise) interface. Three major clusters of inter- and intramolecular interactions of the lysine triad are annotated with interrupted circles/ovals. Lys(K)628 on (ELMO1) primarily engages via strong polar contacts with Met(M)131 and Leu(L)130 on SifA. K626 on (ELMO1) makes an intramolecular contact with D621, a residue within the β5-β6 loop; it is also juxtaposed with K132 and forms a ‘charge-neutralizing’ salt bridge with Asp(D)117 on SifA. K620 on (ELMO1) appears to primarily bind L631 and L638, which are key residue within the β5-β6 loop. **C**. *Top*: An alignment of the sequences of the PH domains of ELMO1 and SKIP is shown, along with secondary structures. Conserved residues are shaded in black; similar residues are boxed. Three lysine residues on ELMO1 that correspond to the structurally resolved contact sites of SKIP for SifA are marked with red boxes. See also **Fig. S3A** for an extended alignment. *Bottom*: A domain map of ELMO1. **D**. An alignment of the β5-β6 loop of ELMO1 showing that the lysine triad highlighted with red box in C is conserved across diverse species. Intra-loop interactions are indicated with interrupted arcs on top. Leu(L) and Asp(D) residues which are engaged in these interactions are also conserved and are highlighted with blue boxes. See also **Fig. S4** for extended alignment. **E**. Panel displays APBS (Adaptive Poisson-Boltzmann Solver)-derived surface electrostatics for an all-side chain model of the ELMO1•SifA co-complex, as visualized using Chimaera. Volume surface coloring was set in the default range of -10 (red), through 0 (white), to +10 (blue) kT/e, where negatively charged surfaces are red (-10 kT/e) and positively charged surfaces are blue (+10 kT/e). In the most energetically favorable orientation, charged residues Lys(K)628 and Lys(K)626 on (ELMO1) bring hydrophobic residues Met(M)131 and Leu(L)130 on SifA into proximity [marked by an oval]. See **Fig. S5** for additional views. **F**. Recombinant WT or K3D mutant His-ELMO1-CT proteins (∼2.5 or 5 μg; Input) were used in pulldown assays with immobilized GST alone or GST-SifA (∼7.5 or 10 μg). Bound ELMO1 was visualized by immunoblotting using an anti-His (ELMO1) antibody. GST proteins are visualized by Ponceau S staining. The K3D mutant, displayed slower electrophoretic mobility compared to the WT ELMO1 protein consistently in both reducing and non-reducing gels, and regardless of whether it was expressed in bacteria as recombinant proteins or expressed in mammalian cells, suggesting it is likely to be due to the introduction of negative charge in the form of 3 aspartates. **G**. Quantification of immunoblots in F. Results are displayed as mean ± S.D (n = 3 independent replicates). Statistical significance was determined using an unpaired t-test. **H**. Equal aliquots (5 μg; Input) of recombinant His-SifA and its mutants (L130D, M131D, and LM-DD) were used in pulldown assays with immobilized GST alone or GST-ELMO1-CT (10 μg). Bound SifA proteins were visualized by immunoblotting using an anti-His antibody. GST proteins are visualized by Ponceau S staining. **I**. Recombinant WT or K3D mutant His-ELMO1-CT proteins were used in pulldown assays with GST or GST-SifA (WT or L130D mutant). Bound ELMO1 was visualized by immunoblotting using an anti-His antibody. GST proteins are visualized by Ponceau S staining. **J**. Quantification of immunoblots in I. Results are displayed as mean ± S.D (n = 3 independent replicates). Statistical significance was determined using an unpaired t-test.

We noted that K620 is reported to be ubiquitinated and the Threonine (T) at 618 is phosphorylated (**Fig. S3B**); none of the lysine residues are reported to be impacted by germline SNPs or somatic mutations in cancers. Most importantly, the β5-β6 loop and the lysine triad within this stretch, are evolutionarily conserved from fish to humans, as well as in the homologous members of the family, ELMO2 and ELMO3 (**Figure 2D**; **Fig. S4**).

To analyze the electrostatics for the model of the SifA•ELMO1 complex, we used APBS (Adaptive Poisson-Boltzmann Solver) (40), a widely accepted software for solving the equations of continuum electrostatics for large biomolecular assemblages. We found that in the most energetically favorable orientation, charged residues Lys(K)628 and Lys(K)626 on ELMO1 bring hydrophobic residues Met(M)131 and Leu(L)130 on SifA into proximity (**Figure 2E**; **Fig. S5**). Because the APBS approach allows us to determine the electrostatic interaction profile as a function of the distance between two molecules, we conclude that the lysines K628 and K626, and potentially other amino acids in the β5-β6 loop are key sites on ELMO1 that engage in electrostatic interactions with L130 and M131 on SifA.

We validated the homology model and the nature of the major interactions [i.e., electrostatic] in the assembly of the complex by generating several structure-rationalized mutants of ELMO1 and SifA. The positively charged lysines on ELMO1 were substituted with negatively charged aspartate residues (D; ELMO1-CT-K3D), expecting that such substitution will disrupt the intermolecular electrostatic attractions and destabilize the SifA•ELMO1 complex. These substitutions were expected to also disrupt intramolecular interactions within the β5-β6 loop (**Figure 2D**) and destabilize the highly conserved loop. The hydrophobic residues on SifA, i.e., L130 and M131, were substituted with an unfavorable negatively charged and polar aspartate residue, either alone (L130D; M131D) or in combination (LM>DD). Binding to SifA was significantly reduced in the case of the K3D ELMO1 mutant (**Figure 2F-G**), and binding to ELMO1 was virtually abolished in the case of all the SifA mutants (**Figure 2H**).

Although both L130 and M131 on SifA were predicted to bind ELMO1, L130 was predicted to be the major contributor (accounting for 13 of the 17 intermolecular contact sites; **Supplemental Information 1**). M131, on the other hand, engaged also in numerous *intra*molecular contacts, which suggests that M131 could be important also for protein conformation. We asked if the strong polar contacts between L130(SifA) and the lysine triad (ELMO1) were critical for the SifA•ELMO1 interaction (**Figure 2B**) and tested their relative contributions without disrupting M131(SifA). Pulldown assays showed that SifA•ELMO1 interactions were partially impaired when L130D-SifA and K3D-ELMO1 substitutions when used alone (**Figure 2I-J**) and virtually lost when the mutants were used concomitantly (**Figure 2I-J**).

These findings provide atomic level insights into the nature and composition of the SifA•ELMO1 complex which is assembled when a pair of hydrophobic residues on SifA binds an evolutionarily conserved polar hot spot on ELMO1-PHD. Strong hydrophobic interactions stabilize the SifA•ELMO1 interface which can be selectively disrupted.

### Disrupting the SifA•ELMO1 interface suppresses Rac1 activity and bacterial invasion

To assess the impact of selective disruption of the SifA•ELMO1 interface in the setting of an infection, we used an unbiased network-based approach. We leveraged a previously published (41) SifA interactome, as determined by proximity-dependent biotin labelling (BioID) and used those interactors as ‘seeds’ for fetching additional interactors to build an integrated SifA(*Salmonella*)•host PPI network (see *Methods* for details). The resultant network (**Fig. S6A**) was perturbed by *in-silico* deletion of either ELMO1 (**Figure 3A, Fig. S6B**) or, more specifically, the SifA•ELMO1 interaction (**Figure 3B**). Both modes of perturbation were analyzed by a differential network analysis (with *vs*. without perturbations) using various network metrices (see legends; **Figure 3C, E**). As one would expect, deletion of ELMO1 impacts many proteins (**Figure 3C**; **Fig. S6C-D**), including those engaged in microbe sensing (BAI1, NOD1, NOD2), membrane trafficking along the phagolysosomal pathway (EEA1, Rab5A, Rab9A and LAMP), multiple Src-family kinases (LYN, YES and HCK), inflammatory cytokines (IL1B, CCL2/MCP1, CXCL12, and TNF), as well as immune cell (B cell) and epithelial (adherens junction) pathways (**Figure 3D**). Findings are consistent with prior published work, implicating ELMO1 in facilitating the recruitment of LAMP1 to SCVs (42), mounting a cytokine response (21, 35), as well as regulating epithelial junctions (43) upon *Salmonella* infection. The proteins found to be impacted based on at least two network metrices (**Figure 3C**) are enriched for Rac1 signaling (Rac1, Rac2, Nckap1 and Dok180) and the endolysosomal pathway (LAMP and RABs). The more refined approach of selective deletion of the SifA•ELMO1 interaction yielded, as expected, a smaller list of proteins that mostly concerned with the DOCK1•RAC signaling axis and Src family kinases, HCK, LYN and YES (**Figure 3E-F**). Because phosphorylation of ELMO1 by Src-family kinases such as Src, Fyn (44), and HCK (45) also converge on Rac1 activity, Rac1 signaling, likely via the ELMO1•DOCK1 axis, emerged as the most important function predicted to be impacted in infected cells.

**Figure 3.**
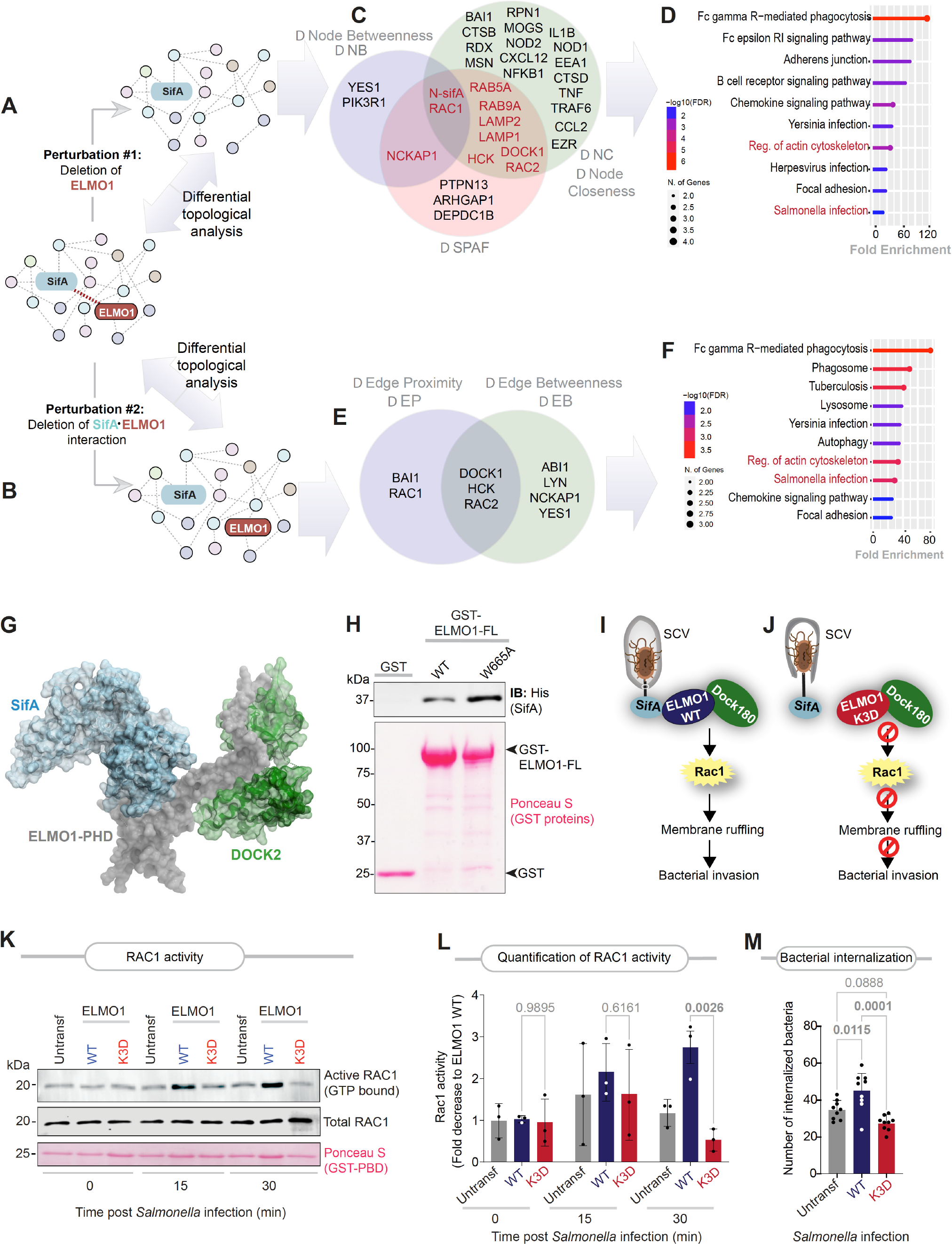
Prediction and validation of the consequences of disruption of the ELMO1•SifA interface. **A-B**. A SifA (*Salmonella*)•human PPI network was constructed using a previously identified BioID-determined interactome of SifA (41) and their connectors were fetched from the human STRING database (54). Network perturbation was carried out *in-silico* either by deletion of ELMO1 (A; node deletion) or selective disruption of the ELMO1•SifA interaction (B; edge deletion). **C-D**. Venn-diagram (C) shows sets of proteins that were impacted by *in-silico* deletion of ELMO1, as determined by three independent metrics of network topological analyses: differential node betweenness (ΔNB), node closeness (ΔNB) and shortest path alteration fraction (ΔSPAF). Lollipop plots (D) indicate the KEGG pathways enriched among the proteins that were identified as impacted based on more than one node-based metric. See **Fig. S6** for a detailed analysis. **E-F**. Venn-diagram (E) shows sets of proteins that were impacted by *in-silico* deletion of the ELMO1•SifA interaction, as determined by two metrics of network topological analyses: differential edge betweenness (ΔEB) and edge proximity (ΔEP). Lollipop plots (F) indicate the KEGG pathways enriched among the proteins that were identified using edge-based metrices. See **Supplemental Information 2** for a detailed list of nodes and edges. **G**. A homology model of a co-complex between SifA (light blue), ELMO1 (gray), and DOCK180 (green), built by overlaying resolved crystal structures of DOCK2•ELMO1 (PDB:3A98)(24), ELMO1 (PDB:2VSZ) and SifA•SKIP (PDB:3CXB). **H**. Recombinant His-SifA (∼5ug) were used in a GST pulldown assay with GST (negative control), and GST-tagged wild-type (WT) and W665A mutants of full length (FL) ELMO1, and bound SifA was visualized by immunoblot using anti-His (SifA) antibody. Equal loading of GST proteins is confirmed by Ponceau S staining. **I-J:** Key steps in SifA•ELMO1•Dock180 co-complex mediated Rac1 signaling (I) and the predicted impact of selectively disrupting SifA•ELMO1 interaction using the K3D mutant (J) are shown. **K-L**. ELMO1-depleted J774 macrophages (Untransfected) transfected with either WT or K3D mutant ELMO1 and subsequently infected with *Salmonella* (MOI 10; at indicated time points post-infection) were assessed for Rac1 activation by pulldown assays using GST-PBD. Immunoblots and Ponceau S-stained GST proteins are shown in K. Quantification of immunoblots in L. Results are displayed as mean ± S.D (n = 3 independent replicates). Statistical significance was determined using one-way ANOVA. **M:** Bar graph represents the number of internalized bacteria by the same cells in K infected with *Salmonella* (MOI 10; 30 min post-infection). Results are displayed as mean ± S.D (n = 3 independent replicates). Statistical significance was determined using one-way ANOVA.

Structure homology models of ternary complexes of SifA•ELMO1•DOCK revealed that although both SifA and DOCK180 bind ELMO1-PHD, they do so via two distinct and non-overlapping interfaces (**Figure 3G**). To experimentally validate this finding, we generated a mutant ELMO1 (W665A) that was previously confirmed by two independent groups to be essential for binding DOCK180 (46, 47) and tested its ability to bind His-SifA in pulldown assays. Both WT and ELMO1-W665A bound SifA to similar extents, indicating that W665 is dispensable for binding SifA (**Figure 3H**). Findings are also consistent with the fact that both SifA and DOCK180 co-immunoprecipitate with ELMO1 (21), and hence, may exist as a ternary complex.

We anticipated that selective disruption of the SifA•ELMO1 interface (using the ELMO1-K3D mutant), while leaving intact the ELMO1•DOCK interface, would interrupt the ELMO1•DOCK•Rac1 signaling axis and reduce bacterial invasion (**Figure 3I-J)**. This was indeed found to be the case as Rac1 signaling was induced during *Salmonella* infection in ELMO1-depleted J774 macrophages reconstituted with WT ELMO1 but found to be significantly blunted when the same macrophages were reconstituted with the K3D mutant ELMO1 (**Figure 3K-L**). Reduced Rac1 activity was also associated with reduced bacterial internalization (**Figure 3M**). Findings demonstrate that one of the major consequences of mutating the polar triad of lysine residues on ELMO1 is reduction in both Rac1 activity and microbial invasion.

### Diverse WxxxE-effectors target the same lysine hotspot on ELMO1

Prior work using ELMO1-*knockout* zebrafish (*E. coli/*MAP (22)) and mouse (*Shigella/*IpgB1 (6) and *Salmonella*/*SifA*) (21) have independently concluded that diverse pathogens trigger host immune responses via *interaction with* ELMO1. We asked if our insights into the nature of the SifA•ELMO1 interface are relevant also to other *WxxxE* motif-containing effectors. As observed previously for SifA (**Figure 1J-K**), *WxxxE*-effectors IpgB1, IpgB2 and Map also directly bound both full length (**Figure 4A***-top*) and the C-terminal PHD containing fragment (**Figure 4A***-bottom*) of ELMO1. More importantly, mutation of the polar lysine triad on ELMO1 reduced the binding of all effectors tested (**Figure 4B-C**), indicating that these effectors require the same hotspot as SifA to engage with ELMO1, presumably via similar hydrophobic contacts. Because the effectors have little to no sequence similarity other than the invariant *WxxxE*-motif (see alignment; **Fig. S7**), and the N-terminal extension is unique to SifA, we were unable to predict the exact nature of the potential hydrophobic contacts on the effectors. We conclude that the newly identified hotspot on ELMO1 represents a point of convergence for numerous *WxxxE*-effectors encoded by diverse pathogens to engage with one host protein. It is noteworthy that each of these *WxxxE*-effectors we tested here, have recently been shown to require ELMO1 for inducing Rac1 signaling (21).

**Figure 4:**
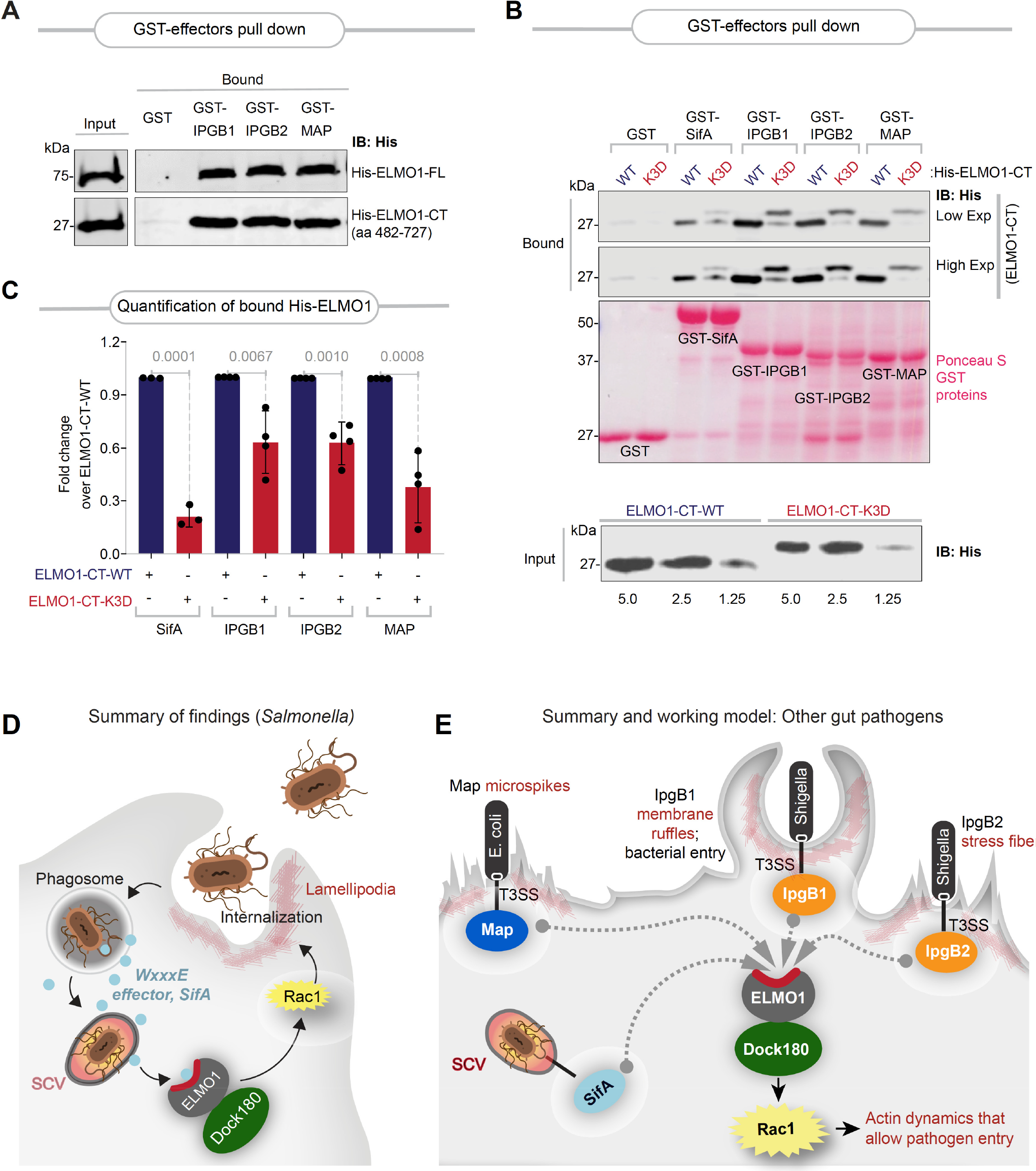
The SifA•ELMO1 interface is a conserved site for engaging diverse WxxxE-effectors. **A**. Recombinant His-tagged CT (aa 482-727) or full-length (FL) ELMO1 proteins (∼3 μg) were used in a pulldown assay with immobilized full-length GST-tagged WxxxE effectors (SifA, IpgB1, IpgB2, Map) or GST alone (negative control). Bound ELMO1 proteins were visualized by immunoblot using anti-His (SifA) antibody. GST proteins are visualized by Ponceau S staining. **B**. Recombinant WT or K3D mutant His-ELMO1-CT proteins (∼3 μg) were used in pulldown assays with immobilized GST-tagged full-length effectors as in A. Bound ELMO1 was visualized by immunoblotting using an anti-His antibody. GST proteins are visualized by Ponceau S staining. **C**. Quantification of immunoblots in B. Results are displayed as mean ± S.D (n = 3 independent replicates). Statistical significance was determined using an unpaired t-test. **D-E**. Schematic (D) summarizes the key findings in this work using *Salmonella* and its WxxxE-motif effector, SifA. Schematic (E) summarizes the key biochemical findings and the working model proposed for how diverse microbes use their *WxxxE* motif-containing virulence factors (effectors) to hijack the host engulfment pathway via a shared mechanism. Although diverse in sequence (see **Fig. S7** for an alignment of WxxxE effectors) and function, they all bind a conserved surface on ELMO1 (highlighted in red). The major consequence of such WxxxE-effector•ELMO1 interaction is enhanced Rac1 activation and bacterial engulfment.

## CONCLUSIONS AND STUDY LIMITATIONS

This work provides an atomic-level insight into a single point of vulnerability within the host engulfment pathway, i.e., a hot spot (lysine triad) on the ELMO1-PHD. This hot spot is exploited by diverse gut pathogens such as *Salmonella* to activate the ELMO1•DOCK180•Rac1 axis, invade host cells and seek refuge within SCVs (see summary of findings; **Figure 4D**). These findings come as a surprise because the bacterial effector proteins that are responsible for such exploitation are diverse, and yet, they all bind the same hot spot on ELMO1 to hijack the Rac1 axis to their advantage (see legend; **Figure 4E**).

Because the *WxxxE*-motif is found in enteric as well as plant pathogens, and within the Toll/interleukin-1 receptor (TIR) modules of both host and pathogen proteins (but notably absent in commensals (21)), it is possible that some host *WxxxE*-proteins may also engage ELMO1 via the same hot spot. If so, the pathogen-encoded *WxxxE*-effectors should competitively block such interactions, exemplifying the phenomenon of molecular mimicry. The structural insights revealed here also warrants the consideration of another form of competitive binding, one in which the same *WxxxE*-effector (SifA) may bind different host proteins (SKIP or ELMO1) using nearly identical interfaces. Because both interactions require the identical residues on SifA, the SifA•ELMO1 and SifA•SKIP interactions must be mutually exclusive. Given the roles of ELMO1 in the regulation of actin dynamics during bacterial entry and SKIP’s role in endosomal tubulation (32) and anterograde movement of endolysosomal compartments (39), we hypothesize that SifA may bind two host proteins sequentially. It may bind ELMO1 first during S*almonella* entry and SCV formation and SKIP later to support cellular processes that help in SCV membrane stabilization, the development of Sifs, and the creation of a favorable environment for the survival and multiplication of *Salmonella* (39, 48). If/how SifA coordinates its interactions with two host proteins, ELMO1 and SKIP, during S*almonella* infection remains unknown; however, the fact that the polar lysine triad on both host proteins is evolutionarily intact from fish to humans suggests that these conserved hot spots on host protein surfaces may have co-evolved with the pathogens as part of a molecular arms race of adaptations. The deleterious impact of structure-rationalized mutants suggests that both the SifA•ELMO1 (shown here) and the SifA•SKIP (published before (26, 32)) interfaces are sensitive to disruption. This is particularly important because hydrophobic interactions that stabilize protein-protein interfaces via a central cluster of hot spot residues are of high therapeutic value because they are amenable to disruption with rationally designed small molecule inhibitors (49).

This study also has a few limitations. For example, how post-translational modifications (PTMs) may impact the *WxxxE*•ELMO1 interface was not evaluated. It is possible that lysine methylation, which ironically was described as a PTM first in *Salmonella* flagellin (50), may impact the interface, as shown in other instances (51). Similarly, phosphorylation at T618 or ubiquitination at K620 may have impacts that were not explored; because the ELMO1•DOCK180 co-complex is known to be regulated by ubiquitination (52), and both proteins are downregulated rapidly upon LPS stimulation(53), we speculate that ubiquitination at K621 may deter *WxxxE*-effector interactions and thereby, serve as a protective strategy of the host during acute infections. Finally, the impact of disrupting the *WxxxE*•ELMO1 interface on host immune responses were not studied here and will require the creation of knock-in K3D mutant models. All these avenues are expected to provide a more complete picture of the consequences of disrupting the *WxxxE*•ELMO1 interface and help formulate strategies to disrupt it for therapeutic purposes.

In conclusion, our studies characterize a polar patch on ELMO1-PHD as a conserved hot spot of host vulnerability—a so-called Achilles heel--which is exploited by multiple pathogens. Because prior studies on ELMO1-*knockout* zebrafish (22) and mice (6, 21) have implicated *WxxxE*•ELMO1 interactions as responsible also for mounting host inflammatory responses, the same polar patch on ELMO1-PHD may also serve as a conserved hot spot of intrusion detection.

## EXPERIMENTAL PROCEDURES

Full experimental procedures can be found in the supporting information.

## RESOURCE AVAILABILITY

### LEAD CONTACT

Further information and requests for resources and reagents should be directed to and will be fulfilled by the lead contact, Pradipta Ghosh.

## MATERIALS AVAILABILITY

This study has generated constructs and cell lines. These materials can only be accessed through proper material transfer agreement following the guidelines of the University of California, San Diego.

## DATA AND CODE AVAILABILITY

All data is available in the main text or the supplementary materials. Original western blot images and microscopy data will be shared by the lead contact upon request. This paper also analyzes an existing, publicly available proteomics dataset and protein structures (listed in the Key Resources Table). Source data for gene ontology analyses are provided with this paper. This paper includes PPI network analysis; a link to the codes is provided (**Key Resources Table**). Any additional information required to reanalyze the data reported in this paper is available from the lead contact upon request.

## CATALOG OF SUPPLORTING MATERIALS

This article contains supporting information that includes.

1. Experimental procedures
2. Supplementary Figures and Legends (7)
3. Supplemental Information (2 Excel Datasheets; uploaded separately)

## ACKNOWLEDGEMENTS

We thank Soumita Das (UC San Diego, CA; currently at University of Massachusetts, Lowell, MA) for numerous reagents and constructs that were used in this work and for helpful discussions. This work was supported by the National Institute of Health (NIH) Grants: R01-AI141630, UG3TR003355, UG3TR002968 and R01-AI55696 (P.G). PG was also supported by the Leona M. and Harry B. Helmsley Charitable Trust and NIH programs. G.D.K. was supported through The American Association of Immunologists (AAI) Intersect Fellowship Program for Computational Scientists and Immunologists. S.R was supported, in part, by the NIH grants (AI118985 and GM117424) and M.A was supported by R01-DK107585 and R01-AI141630. S.R.I was supported by a NIH Diversity Supplement award (3R01DK107585-02S1). We thank Ying Jones (UC San Diego Electron Microscopy Core Facility) for technical and logistical support. The content is solely the responsibility of the authors and does not necessarily represent the official views of the Helmsley Charitable Trust or the National Institutes of Health.

## AUTHOR CONTRIBUTIONS

M. A., and P.G. designed, executed, and analyzed most of the experiments and/or formal analysis in this work. M.A carried out all the biochemical and functional assays. M.A. and G.K. carried out semi-quantitative transmission-electron microscopy (TEM). S. R and P.G, with assistance from A.B, generated and analyzed the sequence and structure homology models, identified structure-rationalized mutants, and carried out the APBS analyses. S. S. created and analyzed the PPI networks. S.R.C. generated the mutant constructs. H.G. and A.A. provided technical assistance for bacterial clearance studies. G.K. and P.G. prepared and/or created display items for data visualization and/or presentation. P.G, with assistance from MA and GK wrote the manuscript and supervised all aspects of this work. All authors reviewed and edited the manuscript and approved its final version.

## REFERENCES

1. Steele-Mortimer, O. (2008) The Salmonella-containing vacuole: moving with the times Curr Opin Microbiol 11, 38–45 10.1016/j.mib.2008.01.002

2. Castanheira, S., andGarcía-Del Portillo, F. (2017) Salmonella Populations inside Host Cells Front Cell Infect Microbiol 7, 432 10.3389/fcimb.2017.00432

3. Lai, Y., Rosenshine, I., Leong, J. M., andFrankel, G. (2013) Intimate host attachment: enteropathogenic and enterohaemorrhagic Escherichia coli Cell Microbiol 15, 1796–1808 10.1111/cmi.12179

4. Smith, K., Humphreys, D., Hume, P. J., andKoronakis, V. (2010) Enteropathogenic Escherichia coli recruits the cellular inositol phosphatase SHIP2 to regulate actin-pedestal formation Cell Host Microbe 7, 13–24 10.1016/j.chom.2009.12.004

5. Watson, J. L., Sanchez-Garrido, J., Goddard, P. J., Torraca, V., Mostowy, S., Shenoy, A. R. et al. (2019) Shigella sonnei O-Antigen Inhibits Internalization, Vacuole Escape, and Inflammasome Activation mBio 10, 10.1128/mBio.02654-19

6. Handa, Y., Suzuki, M., Ohya, K., Iwai, H., Ishijima, N., Koleske, A. J. et al. (2007) Shigella IpgB1 promotes bacterial entry through the ELMO-Dock180 machinery Nat Cell Biol 9, 121–128 10.1038/ncb1526

7. Finlay, B. B., andCossart, P. (1997) Exploitation of mammalian host cell functions by bacterial pathogens Science 276, 718–725 10.1126/science.276.5313.718

8. Galán, J. E., andCollmer, A. (1999) Type III secretion machines: bacterial devices for protein delivery into host cells Science 284, 1322–1328 10.1126/science.284.5418.1322

9. Cossart, P., andSansonetti, P. J. (2004) Bacterial invasion: the paradigms of enteroinvasive pathogens Science 304, 242–248 10.1126/science.1090124

10. Connor, M. G., Pulsifer, A. R., Chung, D., Rouchka, E. C., Ceresa, B. K., andLawrenz, M. B. (2018) Yersinia pestis Targets the Host Endosome Recycling Pathway during the Biogenesis of the Yersinia-Containing Vacuole To Avoid Killing by Macrophages mBio 9, 10.1128/mBio.01800-17

11. Watson, R. O., andGalán, J. E. (2008) Campylobacter jejuni survives within epithelial cells by avoiding delivery to lysosomes PLoS Pathog 4, e14 10.1371/journal.ppat.0040014

12. Coburn, B., Sekirov, I., andFinlay, B. B. (2007) Type III secretion systems and disease Clin Microbiol Rev 20, 535–549 10.1128/cmr.00013-07

13. Orchard, R. C., Kittisopikul, M., Altschuler, S. J., Wu, L. F., Süel, G. M., andAlto, N. M. (2012) Identification of F-actin as the dynamic hub in a microbial-induced GTPase polarity circuit Cell 148, 803–815 10.1016/j.cell.2011.11.063

14. Alto, N. M., Shao, F., Lazar, C. S., Brost, R. L., Chua, G., Mattoo, S. et al. (2006) Identification of a bacterial type III effector family with G protein mimicry functions Cell 124, 133–145 10.1016/j.cell.2005.10.031

15. Bulgin, R., Raymond, B., Garnett, J. A., Frankel, G., Crepin, V. F., Berger, C. N. et al. (2010) Bacterial guanine nucleotide exchange factors SopE-like and WxxxE effectors Infect Immun 78, 1417–1425 10.1128/iai.01250-09

16. Hardt, W. D., Chen, L. M., Schuebel, K. E., Bustelo, X. R., andGalán, J. E. (1998) S. typhimurium encodes an activator of Rho GTPases that induces membrane ruffling and nuclear responses in host cells Cell 93, 815–826 10.1016/s0092-8674(00)81442-7

17. Ohya, K., Handa, Y., Ogawa, M., Suzuki, M., andSasakawa, C. (2005) IpgB1 is a novel Shigella effector protein involved in bacterial invasion of host cells. Its activity to promote membrane ruffling via Rac1 and Cdc42 activation J Biol Chem 280, 24022–24034 10.1074/jbc.M502509200

18. Bulgin, R. R., Arbeloa, A., Chung, J. C., andFrankel, G. (2009) EspT triggers formation of lamellipodia and membrane ruffles through activation of Rac-1 and Cdc42 Cell Microbiol 11, 217–229 10.1111/j.1462-5822.2008.01248.x

19. Arbeloa, A., Bulgin, R. R., MacKenzie, G., Shaw, R. K., Pallen, M. J., Crepin, V. F. et al. (2008) Subversion of actin dynamics by EspM effectors of attaching and effacing bacterial pathogens Cell Microbiol 10, 1429–1441 10.1111/j.1462-5822.2008.01136.x

20. Kenny, B., Ellis, S., Leard, A. D., Warawa, J., Mellor, H., andJepson, M. A. (2002) Co-ordinate regulation of distinct host cell signalling pathways by multifunctional enteropathogenic Escherichia coli effector molecules Mol Microbiol 44, 1095–1107 10.1046/j.1365-2958.2002.02952.x

21. Sayed, I. M., Ibeawuchi, S. R., Lie, D., Anandachar, M. S., Pranadinata, R., Raffatellu, M. et al. (2021) The interaction of enteric bacterial effectors with the host engulfment pathway control innate immune responses Gut Microbes 13, 1991776 10.1080/19490976.2021.1991776

22. Xue, R., Wang, Y., Wang, T., Lyu, M., Mo, G., Fan, X. et al. (2021) Functional Verification of Novel ELMO1 Variants by Live Imaging in Zebrafish Front Cell Dev Biol 9, 723804 10.3389/fcell.2021.723804

23. Grimsley, C. M., Kinchen, J. M., Tosello-Trampont, A. C., Brugnera, E., Haney, L. B., Lu, M. et al. (2004) Dock180 and ELMO1 proteins cooperate to promote evolutionarily conserved Rac-dependent cell migration J Biol Chem 279, 6087–6097 10.1074/jbc.M307087200

24. Hanawa-Suetsugu, K., Kukimoto-Niino, M., Mishima-Tsumagari, C., Akasaka, R., Ohsawa, N., Sekine, S. et al. (2012) Structural basis for mutual relief of the Rac guanine nucleotide exchange factor DOCK2 and its partner ELMO1 from their autoinhibited forms Proc Natl Acad Sci U S A 109, 3305–3310 10.1073/pnas.1113512109

25. Kukimoto-Niino, M., Ihara, K., Murayama, K., andShirouzu, M. (2021) Structural insights into the small GTPase specificity of the DOCK guanine nucleotide exchange factors Curr Opin Struct Biol 71, 249–258 10.1016/j.sbi.2021.08.001

26. Diacovich, L., Dumont, A., Lafitte, D., Soprano, E., Guilhon, A. A., Bignon, C. et al. (2009) Interaction between the SifA virulence factor and its host target SKIP is essential for Salmonella pathogenesis J Biol Chem 284, 33151–33160 10.1074/jbc.M109.034975

27. Lin, Q., Yang, W., Baird, D., Feng, Q., andCerione, R. A. (2006) Identification of a DOCK180-related guanine nucleotide exchange factor that is capable of mediating a positive feedback activation of Cdc42 J Biol Chem 281, 35253–35262 10.1074/jbc.M606248200

28. Toret, C. P., Collins, C., andNelson, W. J. (2014) An Elmo-Dock complex locally controls Rho GTPases and actin remodeling during cadherin-mediated adhesion J Cell Biol 207, 577–587 10.1083/jcb.201406135

29. Achi, S. C., Karimilangi, S., Lie, D., Sayed, I. M., andDas, S. (2023) The WxxxE proteins in microbial pathogenesis Crit Rev Microbiol 49, 197–213 10.1080/1040841x.2022.2046546

30. Huang, Z., Sutton, S. E., Wallenfang, A. J., Orchard, R. C., Wu, X., Feng, Y. et al. (2009) Structural insights into host GTPase isoform selection by a family of bacterial GEF mimics Nat Struct Mol Biol 16, 853–860 10.1038/nsmb.1647

31. Klink, B. U., Barden, S., Heidler, T. V., Borchers, C., Ladwein, M., Stradal, T. E. et al. (2010) Structure of Shigella IpgB2 in complex with human RhoA: implications for the mechanism of bacterial guanine nucleotide exchange factor mimicry J Biol Chem 285, 17197–17208 10.1074/jbc.M110.107953

32. Ohlson, M. B., Huang, Z., Alto, N. M., Blanc, M. P., Dixon, J. E., Chai, J. et al. (2008) Structure and function of Salmonella SifA indicate that its interactions with SKIP, SseJ, and RhoA family GTPases induce endosomal tubulation Cell Host Microbe 4, 434–446 10.1016/j.chom.2008.08.012

33. Arbeloa, A., Garnett, J., Lillington, J., Bulgin, R. R., Berger, C. N., Lea, S. M. et al. (2010) EspM2 is a RhoA guanine nucleotide exchange factor Cell Microbiol 12, 654–664 10.1111/j.1462-5822.2009.01423.x

34. Raymond, B., Crepin, V. F., Collins, J. W., andFrankel, G. (2011) The WxxxE effector EspT triggers expression of immune mediators in an Erk/JNK and NF-κB-dependent manner Cell Microbiol 13, 1881–1893 10.1111/j.1462-5822.2011.01666.x

35. Das, S., Sarkar, A., Choudhury, S. S., Owen, K. A., Castillo, V., Fox, S. et al. (2015) ELMO1 has an essential role in the internalization of Salmonella Typhimurium into enteric macrophages that impacts disease outcome Cell Mol Gastroenterol Hepatol 1, 311–324 10.1016/j.jcmgh.2015.02.003

36. Ruiz-Albert, J., Yu, X. J., Beuzón, C. R., Blakey, A. N., Galyov, E. E., andHolden, D. W. (2002) Complementary activities of SseJ and SifA regulate dynamics of the Salmonella typhimurium vacuolar membrane Mol Microbiol 44, 645–661 10.1046/j.1365-2958.2002.02912.x

37. Beuzón, C. R., Méresse, S., Unsworth, K. E., Ruíz-Albert, J., Garvis, S., Waterman, S. R. et al. (2000) Salmonella maintains the integrity of its intracellular vacuole through the action of SifA Embo j 19, 3235–3249 10.1093/emboj/19.13.3235

38. Arpaia, N., Godec, J., Lau, L., Sivick, K. E., McLaughlin, L. M., Jones, M. B. et al. (2011) TLR signaling is required for Salmonella typhimurium virulence Cell 144, 675–688 10.1016/j.cell.2011.01.031

39. Dumont, A., Boucrot, E., Drevensek, S., Daire, V., Gorvel, J. P., Poüs, C. et al. (2010) SKIP, the host target of the Salmonella virulence factor SifA, promotes kinesin-1-dependent vacuolar membrane exchanges Traffic 11, 899–911 10.1111/j.1600-0854.2010.01069.x

40. Jurrus, E., Engel, D., Star, K., Monson, K., Brandi, J., Felberg, L. E. et al. (2018) Improvements to the APBS biomolecular solvation software suite Protein Sci 27, 112–128 10.1002/pro.3280

41. D’Costa, V. M., Coyaud, E., Boddy, K. C., Laurent, E. M. N., St-Germain, J., Li, T. et al. (2019) BioID screen of Salmonella type 3 secreted effectors reveals host factors involved in vacuole positioning and stability during infection Nat Microbiol 4, 2511–2522 10.1038/s41564-019-0580-9

42. Sarkar, A., Tindle, C., Pranadinata, R. F., Reed, S., Eckmann, L., Stappenbeck, T. S. et al. (2017) ELMO1 Regulates Autophagy Induction and Bacterial Clearance During Enteric Infection J Infect Dis 216, 1655–1666 10.1093/infdis/jix528

43. Sayed, I. M., Suarez, K., Lim, E., Singh, S., Pereira, M., Ibeawuchi, S. R. et al. (2020) Host engulfment pathway controls inflammation in inflammatory bowel disease Febs j 287, 3967–3988 10.1111/febs.15236

44. Makino, Y., Tsuda, M., Ohba, Y., Nishihara, H., Sawa, H., Nagashima, K. et al. (2015) Tyr724 phosphorylation of ELMO1 by Src is involved in cell spreading and migration via Rac1 activation Cell Commun Signal 13, 35 10.1186/s12964-015-0113-y

45. Yokoyama, N., deBakker, C. D., Zappacosta, F., Huddleston, M. J., Annan, R. S., Ravichandran, K. S. et al. (2005) Identification of tyrosine residues on ELMO1 that are phosphorylated by the Src-family kinase Hck Biochemistry 44, 8841–8849 10.1021/bi0500832

46. Kukimoto-Niino, M., Katsura, K., Kaushik, R., Ehara, H., Yokoyama, T., Uchikubo-Kamo, T. et al. (2021) Cryo-EM structure of the human ELMO1-DOCK5-Rac1 complex Sci Adv 7, 10.1126/sciadv.abg3147

47. Lu, M., Kinchen, J. M., Rossman, K. L., Grimsley, C., deBakker, C., Brugnera, E. et al. (2004) PH domain of ELMO functions in trans to regulate Rac activation via Dock180 Nat Struct Mol Biol 11, 756–762 10.1038/nsmb800

48. Jackson, L. K., Nawabi, P., Hentea, C., Roark, E. A., andHaldar, K. (2008) The Salmonella virulence protein SifA is a G protein antagonist Proc Natl Acad Sci U S A 105, 14141–14146 10.1073/pnas.0801872105

49. Li, Y., Huang, Y., Swaminathan, C. P., Smith-Gill, S. J., andMariuzza, R. A. (2005) Magnitude of the hydrophobic effect at central versus peripheral sites in protein-protein interfaces Structure 13, 297–307 10.1016/j.str.2004.12.012

50. Ambler, R. P., andRees, M. W. (1959) Epsilon-N-Methyl-lysine in bacterial flagellar protein Nature 184, 56–57 10.1038/184056b0

51. Huang, J., Sengupta, R., Espejo, A. B., Lee, M. G., Dorsey, J. A., Richter, M. et al. (2007) p53 is regulated by the lysine demethylase LSD1 Nature 449, 105–108 10.1038/nature06092

52. Makino, Y., Tsuda, M., Ichihara, S., Watanabe, T., Sakai, M., Sawa, H. et al. (2006) Elmo1 inhibits ubiquitylation of Dock180 J Cell Sci 119, 923–932 10.1242/jcs.02797

53. Song, J. H., Mascarenhas, J. B., Sammani, S., Kempf, C. L., Cai, H., Camp, S. M. et al. (2022) TLR4 activation induces inflammatory vascular permeability via Dock1 targeting and NOX4 upregulation Biochim Biophys Acta Mol Basis Dis 1868, 166562 10.1016/j.bbadis.2022.166562

54. Szklarczyk, D., Gable, A. L., Nastou, K. C., Lyon, D., Kirsch, R., Pyysalo, S. et al. (2021) The STRING database in 2021: customizable protein-protein networks, and functional characterization of user-uploaded gene/measurement sets Nucleic Acids Res 49, D605–D612 10.1093/nar/gkaa1074

