## Supplementary Online Materials for "Diverse gut pathogens exploit the host engulfment pathway via a conserved mechanism"

**AFFILIATIONS:**
[1]*Department of Cellular and Molecular Medicine, University of California San Diego.*
[2]*Department of Pathology, University of California San Diego.*
[3]*Department of Medicine, University of California San Diego.*

**RUNNING TITLE:** Gut pathogens exploit host's engulfment pathway

***CORRESPONDING AUTHORS:**

### CATALOG OF SUPPORTING MATERIALS

1. *Key Resources and Detailed Experimental procedures*
2. *Supplementary Figures and Legends (7)*
3. *Supplemental Information (2 Excel Datasheets; uploaded separately)*

### Experimental procedures

#### TABLE OF KEY RESOURCES

| REAGENT or RESOURCE | SOURCE | IDENTIFIER |
|---|---|---|
| Antibodies | | |
| Mouse monoclonal anti-poly-His | Sigma-Aldrich | Cat# H1029 |
| Mouse monoclonal anti-FLAG | Cell Signaling | Cat# 8146S |
| Mouse monoclonal ELMO1 antibody | Santa Cruz Biotechnology | Cat# sc-271519 |
| IRDye 800CW Goat anti-Mouse IgG Secondary | LI-COR Biosciences | Cat# 926-32210 |
| IRDye 680RD Goat anti-Rabbit IgG Secondary | LI-COR Biosciences | Cat# 926-68071 |
| **Biological samples** | | |
| HEK 293 | ATCC | CRL-1573 |
| sh Control J774 and sh ELMO1 J774 murine macrophage cell lines | Sayed et al., 2021(1) | n/a |
| *Salmonella enteric serovar Typhimurium* (strain SL1344) | ATCC | Cat#700720 |
| *SifA* mutant strain | Obtained from Olivia Steele-Mortimer (NIH/NIAID) | n/a |
| **Chemicals, peptides** | | |
| HisPur™ Cobalt Resin | Thermo Scientific | Cat# 89964 |
| Glutathione Sepharose® 4B | Sigma-Aldrich | Cat# GE17-0756-04 |
| PVDF Transfer Membrane, 0.45mM | Thermo Scientific | Cat# 88518 |
| Polyethylenimine (PEI) | Polysciences | Cat# Inc 23966 |
| Protease inhibitor cocktail | Roche | Cat# 11 873 580 001 |
| **Critical commercial assay kits** | | |
| Rac1 Pull-Down Activation Assay Biochem Kit (Bead Pull-Down Format) | Cytoskeleton.Inc. | Cat# BK035-S |
| **Plasmids and recombinant proteins** | | |
| pGEX-6P-2 (GST) | GE Healthcare | Cat# 28-9546-50 |
| GST ELMO1 full-length (FL) | Sayed et al., 2021(1) | n/a |
| GST ELMO1 CT (aa 482-727) | *This paper* | n/a |
| GST SifA WT | Sayed et al., 2021(1) | n/a |
| GST SifA L130D | *This paper* | n/a |
| GST-IPGB1 | *This paper* | Cloned from FLAG-IPGB1 provided by Neil Alto, UT Dallas |
| GST-IPGB2 | *This paper* | Cloned from FLAG-IPGB2 provided by Neil Alto, UT Dallas |
| GST- MAP | *This paper* | Cloned from FLAG-Map provided by Neil Alto, UT Dallas |
| pET-28a (His) | Novagene | Cat# 69864 |
| His ELMO1 full-length (FL) | *This paper* | n/a |
| His ELMO1 CT (aa 482-727) | *This paper* | n/a |
| His ELMO1 K620D/K626D/K628D | *This paper* | n/a |
| His SifA WT | *This paper* | n/a |
| His SifA L130D | *This paper* | n/a |

| | | |
|---|---|---|
| His SifA M131D | *This paper* | n/a |
| His SifA L130D/M131D | *This paper* | n/a |
| FLAG ELMO1 (WT) | Sayed et al., 2021(1) | n/a |
| FLAG ELMO1 (K3D) | *This paper* | K620D; K626D; K628D |
| **Mutagenesis Primers** | **Forward primer** | **Reverse primer** |
| Mouse ELMO1 K620D/K626D/K628D | 5'-tgttttgtttaagggcaccatcctcatccatatgagggcagtcctttcccgtc-3'<br><br>5'-gacgggaaaggactgccctcatatggatgaggatggtgcccttaaacaaaaca-3'<br><br>5'-caaagcagtggtgacgggagatgactgccctcatatgaaag-3' | 5'-ctttcatatgagggcagtcatctcccgtcaccactgctttg-3' |
| SifA L130D | 5'-ccgggcgatctttcatatcaaaataaagattccccatgacttcacagct-3' | 5'-agctgtgaagtcatggggaatctttattttgatatgaaagatcgcccgg-3' |
| SifA M131D | 5'-ttaaaatatccgggcgatctttatcatcaaaataaagattccccatgacttcacagc-3' | 5'-gctgtgaagtcatggggaatctttattttgatgataaagatcgcccggatattttaa-3' |
| **Cloning Primers** | **Forward primer (5'→3')** | **Reverse primer (5'→3')** |
| Elmo1FL | | |
| Elmo1 CT (BamHI/Not1) | cgcggatccatgcaggtggtgaaggagcagg | gcgacgcggccgctgttacagtcatagacaaagtcatagttgctaggttcct |
| IPGB1 (KpnI/BamHI) | catggtaccatgcaaattctaaacaaaatacttccacaggt | cgcggatccttaatttgtattgctttgacggtatacagcctt |
| IPGB2 (KpnI/BamHI) | catggtaccatgcttggaacatcttttaataattttggaatcagt | cgcggatcctcagaaaggcgattctaaatttgtaatatagtcac |
| MAP (EcoRI/KpnI) | ccggaattccatgtttagtccaatgacaatggcagg | catggtaccctacaatcgggtatcctgtacatgctc |
| **Software and algorithms** | | |
| ImageStudio Lite | LI-COR | https://www.licor.com/bio/image-studio-lite/ |
| Prism | GraphPad | Home - GraphPad |
| PyMOL Molecular Graphics System | | https://pymol.org/2 |
| ShinyGO v0.76 | | http://bioinformatics.sdstate.edu/go/ |
| Codes for the construction and analysis of the PPI networks | | https://github.com/sinha7290/ELMO |
| **Publicly deposited structures and/or datasets** | | |
| ELMO1 (PH domain) | PDB:2VSZ | https://www.rcsb.org/structure/2VSZ |
| SKIP-PH bound to SifA | PDB:3CXB | https://www.rcsb.org/structure/3CXB |
| DOCK2-bound to ELMO1 | PDB:3A98 | https://www.rcsb.org/structure/3A98 |
| Proximity ligation assay (BioID) | SifA interactome | D'Costa VM et al.,(2) |

### EXPERIMENTAL MODEL AND SUBJECT DETAILS

J774 macrophage and human (HEK) cell lines were used in various assays. Bacterial strains were used to infect macrophages in various experiments. This study does not involve the use of human subjects or animals.

### METHOD DETAILS

#### *Plasmid constructs and mutagenesis.*

ELMO1 and SifA (from SL1344) were amplified by PCR and cloned into pET-28a (+) and pGEX-6P-2 plasmid vectors to generate His- and GST-tagged proteins, respectively. ELMO1 CT (aa 482-727) was amplified by PCR from human ELMO1 cDNA (Clone IRAUp969C0316D, SourceBioscience Inc) and subcloned into pET-28a (+) and pGEX-6P-2 plasmid vectors using BamHI and NotI restriction sites. Effector proteins IPGB1, IPGB2 and MAP were amplified from FLAG-tagged effector constructs (obtained from Neil Alto, UT Dallas) and cloned into pGEX-6P-2 plasmid vectors. ELMO1 and SifA mutations were generated by site directed mutagenesis using the Quick-Change II Site-Directed Mutagenesis kit (Agilent Technologies) and specific primers (**table of key resources**) according to manufacturer's protocol. All plasmid constructs and mutagenesis were verified by sequencing and protein expression was verified by western blot analysis.

#### *Bacteria and bacterial culture*

*Salmonella* enterica serovar *Typhimurium* strain SL1344, were obtained from the American Type Culture Collection (ATCC) (Manassas, VA, USA), and SL1344 Δ*SifA* mutant strain was obtained from Olivia Steele Mortimer (NIH/NIAID) (3, 4) . The bacterial strains were propagated as described previously (5, 6). Briefly, a single colony was inoculated into LB broth and grown for 8 h under aerobic conditions and then under oxygen-limiting conditions. For SL1344 Δ*SifA* mutant strain, streptomycin with final conc (100 µg/mL) was added to LB broth. Cells were infected with a multiplicity of infection (moi) of 10.

#### *Cell culture and transfection*

HEK cells were obtained from American Type Culture Collection, (Manassas, VA, USA). Control and ELMO1 depleted shRNA J774 cells were generated as previously described (7). Cells were maintained in high glucose DMEM (Life Technologies) containing 10% fetal bovine serum and 100 U/ml penicillin and streptomycin at 37$^{o}$C in a 5% CO2 incubator. Cells were sub-cultured 24 h prior to transfection. Transfections of plasmids were performed using Lipofectamine 2000 (Invitrogen) according to manufacturer's protocol.

#### *Transmission electron microscopy (TEM)*

Control (shControl) and ELMO1-depleted (shELMO1) J477 macrophages were infected with *Salmonella* enterica serovar *Typhimurium* strain SL1344 (SL) and its mutant strain *SifA* [MOI 30] and the infected pellets were fixed with Glutaraldehyde in 0.1 M Sodium Cacodylate Buffer, (pH 7.4) followed by post-fixation with 1% OsO4 in 0.1 M cacodylate buffer for 1 h on ice followed by staining with 2% uranyl acetate for 1 h and dehydrated in ethanol

(50-100%) on ice. The cells were washed for 10 min each once with 100% ethanol and two times with acetone and embedded with Durcupan. A Leica UCT ultramicrotome was used to section 60 nm cuts and picked up on 300 mesh copper grids. Sections were post-stained first with 2% uranyl acetate (5 min) and Sato's lead stain (1 min) prior to visualization using a JEOL-1400plus equipped with a bottom-mount Gatan (4k x 4k) camera.

***Protein expression and purification***

Recombinant His and GST-tagged proteins were expressed in Escherichia coli strain BL21 (DE3) and purified as previously described (1, 2). Briefly, proteins in bacterial culture were induced with IPTG (1 mM) overnight incubation at 25°C. Centrifuged and cell pellet were then lysed in either GST lysis buffer containing 25 mM Tris-HCl (pH 7.4) 20 mM NaCl, 1 mM EDTA, 20% (v/v) glycerol, 1% (v/v) Triton X-100, protease inhibitor cocktail or His lysis buffer containing 50 mM $NaH_2PO_4$ (pH 7.4), 300 mM NaCl, 10 mM imidazole, 1% (v/v) Triton-X-100, protease inhibitor cocktail and then sonicated briefly to ensure better lysis. Cell lysate were centrifugation at 12,000 x g at 4°C for 30 mins to remove cell debris. Cleared cell lysate was then affinity purified using either glutathione-Sepharose 4B beads or HisPur Cobalt Resin, followed by elution and overnight dialysis in PBS. Proteins were aliquoted and stored at -80°C prior to use.

***GST pulldown assays***

Recombinant purified GST-tagged proteins were immobilized onto glutathione-Sepharose beads in a binding buffer containing 50 mM Tris-HCl (pH 7.4), 100 mM NaCl, 0.4% (v/v) Nonidet P-40, 10 mM $MgCl_2$, 5 mM EDTA, 2 mM DTT for 60 min at 4°C. GST-tagged protein bound beads were washed and incubated with purified His-tagged proteins in a binding buffer for 4 h at 4°C. Protein bound complexes on a beads were washed four times with phosphate wash buffer containing 4.3 mM $Na_2HPO_4$, 1.4 mM $KH_2PO_4$ (pH 7.4), 137 mM NaCl, 2.7 mM KCl, 0.1% (v/v) Tween-20, 10 mM $MgCl_2$, 5 mM EDTA, 2 mM DTT, 0.5 mM sodium orthovanadate and eluted by boiling beads in reducing sample buffer containing 5% SDS, 156 mM Tris-Base, 25% glycerol, 0.025% bromophenol blue, 25% β-mercaptoethanol. Bound proteins and lysates were fractionated by SDS/PAGE gel and analyzed by immunoblot.

***Quantitative immunoblotting***

For immunoblotting, protein samples were separated by 10% SDS-PAGE and transferred to polyvinylidene fluoride (PVDF) membranes. Membranes were blocked with 5% non-fat milk in Tris-buffered saline with 0.1% Tween® 20 detergent (1X TBST). The membrane was stained with Ponceau S to visualize bait proteins (GST), washed, blocked (5% milk), and incubated with primary antibody solutions overnight at 4°C (unless otherwise specified, a 1:500 dilution for primary antibody was used). Washed blots were then incubated with infrared secondary antibodies (see Key Resource Table) for 1 h at room temperature. Dual color infrared imaging and quantifications of immunoblots were performed using a Li-Cor Odyssey imaging system and analyzed using the Image StudioLite software as per manufacturer's instructions.

***Homology modelling and protein docking***

For the structure-guided mutagenesis studies, we took into consideration the following three previously resolved structures, all accessed through the Protein Data Bank (PDB; https://www.rcsb.org/)-- (i) PDB:3CXB, a 2.60 Å resolved crystal structure of the PH domain of SKIP (3CXB, golden brown) in complex with full length SifA (3CXB, cyan; (ii) PDB:2VSZ, a 2.30 Å resolved crystal structure of the C-terminal 200 aa (containing the PH domain) of ELMO1 in complex with the N-terminal 200 aa of DOCK180; (iii) PDB:3A98, a 2.10 N-terminal 177 aa fragment of DOCK2 and the C-terminal 196 aa fragment (containing the PH domain) of ELMO1. First, the PH domain of SKIP (PDB:3CXB; template) was superimposed to the PH-domain of ELMO1 (PDB:2VSZ). The performed analysis of the PH domains revealed them to be quite similar in structure SKIP and ELMO1 (RMSD of 1.528). Subsequently, the PH domain was used for molecular docking with SifA. Protein docking and the prediction of the contact sites at the ELMO1•SifA interface was performed by using HADDOCK (High Ambiguity Driven protein−protein Docking (8)) web server and the final representation was using Pymol visualization tool (https://pymol.org/2/; The PyMOL Molecular Graphics System. Delano Scientific, San Carlos). HADDOCK is a popular docking program that takes a data-driven approach to docking, with support for a wide range of experimental data (8). HADDOCK generated **130** structures in **15** cluster(s), which represents **65 %** of the water-refined models. The HADDOCK scoring function consists of a linear combination of various energies and buried surface area (HADDOCK2.2 scoring function). Based on such HADDOCK scoring, the models in cluster #3 were most optimal based on the Z-score which indicates how many standard deviations from the average this cluster is located in terms of score (the more negative the better). For cluster 3 we got a Z score of -1.5 and HADDOCK score of -59.7 (**Fig. S1**). The best fit co-complex model was used for predicting the properties and nature of the protein-protein binding interface. A complete list of contacts is provided in **Supplemental Information 1** as an excel datasheet. For calculating the electrostatic surface potential (kT/e) for the ELMO1•SifA co-complex we used adaptive Poisson-Boltzmann solver (APBS), an NIH-supported software (9)[https://www.poissonboltzmann.org/] for the analysis of electrostatic forces for large bimolecular assemblages. The APBS derived surface potential of the ELMO:SifA complex was visualized using Chimaera UCSF software (10). Amino acids were color-coded (red negatively charged; blue positively charged) based on the values of APBS scale in a range of -10 to +10 kT/e as default settings.

***Construction, perturbation, and analysis of the Protein-Protein Interaction Network***

A SifA (*Salmonella*)•human interaction network was constructed using a previously identified BioID-determined interactome of SifA (11) and their connectors, fetched from the human STRING database (12, 13). To avoid false positive interactions, a Stepminer-derived cutoff of combined interaction score was selected (12, 13). A complete list of edges is provided in **Supplemental Information 1**.

Once constructed, the network was perturbed using two approaches. First, an *in-silico* deletion of ELMO1 was performed, and each node within this perturbed network (without ELMO1) was assessed for the following network metrics: (i) Betweenness centrality, (ii) Closeness centrality, and (iii) Shortest Path Alteration Fraction (SPAF) (14). Difference in Z score, $\Delta Z$, of each node, N, for each of the above mentioned topological metrices was calculated. Proteins with $\Delta Z > 0.1$ were identified as those that were impacted due to the deletion of ELMO1.

As a second approach, an *in-silico* deletion of ELMO1•SifA interaction(edge) was performed, and each edge within this perturbed network (without ELMO1•SifA edge) was assessed for two independent metrices: (i) Edge betweenness centrality and (ii) Edge Proximity (15). Edges with $\Delta Z > 0.1$ were identified as those that were impacted due to the selective deletion of the ELMO1•SifA interaction.

A group of proteins that are impacted, as determined by at least more than one topological node-based metrices, or either of the two edge-based metrics, were used for KEGG enrichment analysis (using ShinyGO v0.76; http://bioinformatics.sdstate.edu/go/). All the codes for network analysis are available at https://github.com/sinha7290/ELMO.

***Measurement of Rac1 activity***

ELMO1-depleted murine J774 macrophage cell lines were transfected with ELMO1 WT and ELMO1 K3D plasmids and infected with *Salmonella enterica* serovar Typhimurium for 15 and 30 min (MOI 1:50) and Rac1 activity was measured by pull-down assay using a commercial kit. First, infected cells were lysed with the lysis buffer provided in the kit. Equal aliquots of cell lysates (~ 800 µg total cell protein) were added to 10 µg (10 µl) of immobilized GST-PBD (glutathione *S*-transferase with p21-binding domain of Pak1) (a.k.a., PAK-PBD beads) and incubated at 4°C on a rotator for 1 h. PAK-PBD beads were recovered by centrifugation at 5,000 x g at 4°C. Unbound lysates were removed and the beads were washed once with 500 µl Wash Buffer prior to eluting the proteins with 20 µl of 2x Laemmli sample buffer and boiling for 5 min at 100°C. Input lysates and eluates from the PAK-PBD beads were resolved on a 12.5% SDS-PAGE, and transferred for 45 min, followed by quantitative immunoblotting.

***Bacterial internalization assays***

Approximately $0.5*10^6$ ELMO1-depleted murine macrophage cells were seeded into 12-well culture dishes ~12 h before transfection with either FLAG-ELMO1 WT or FLAG-ELMO1 K3D. After 48 hours of transfection, cells were infected with SL at a MOI of 1:50 for 30 min in antibiotic free media. The cells were then washed and incubated with gentamicin (500 mg/ml) for 90 min to kill the extracellular bacteria. Subsequently, the cells were lysed in 1% Triton-X100, and the lysates were serially diluted and plated directly onto Luria-Bertani agar plates. The total colony-forming units (CFUs) were enumerated the next day after overnight incubation of the plates at 37°C.

***Image Processing***

All images were processed on ImageJ software (NIH) or iStudio (LiCOR) software and assembled into figure panels using Photoshop and Illustrator (Adobe Creative Cloud). All graphs were generated using GraphPad Prism v9.3.1.

### QUANTIFICATION AND STATISTICAL ANALYSIS

*Statistical Analysis and Replicates*

All experiments were repeated at least three times, and results were presented as average ± S.E.M. Statistical significance was assessed using either t-test or one-way analysis of variance (ANOVA) including a Tukey's test for multiple comparisons. Actual *p* values are displayed.

### Supplementary Figures and Legends

A

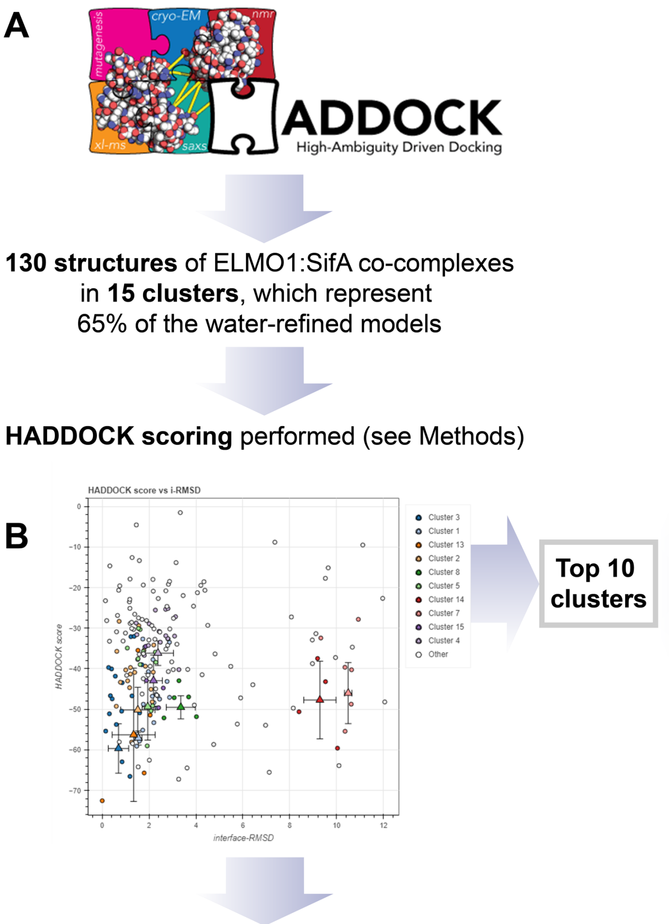

C

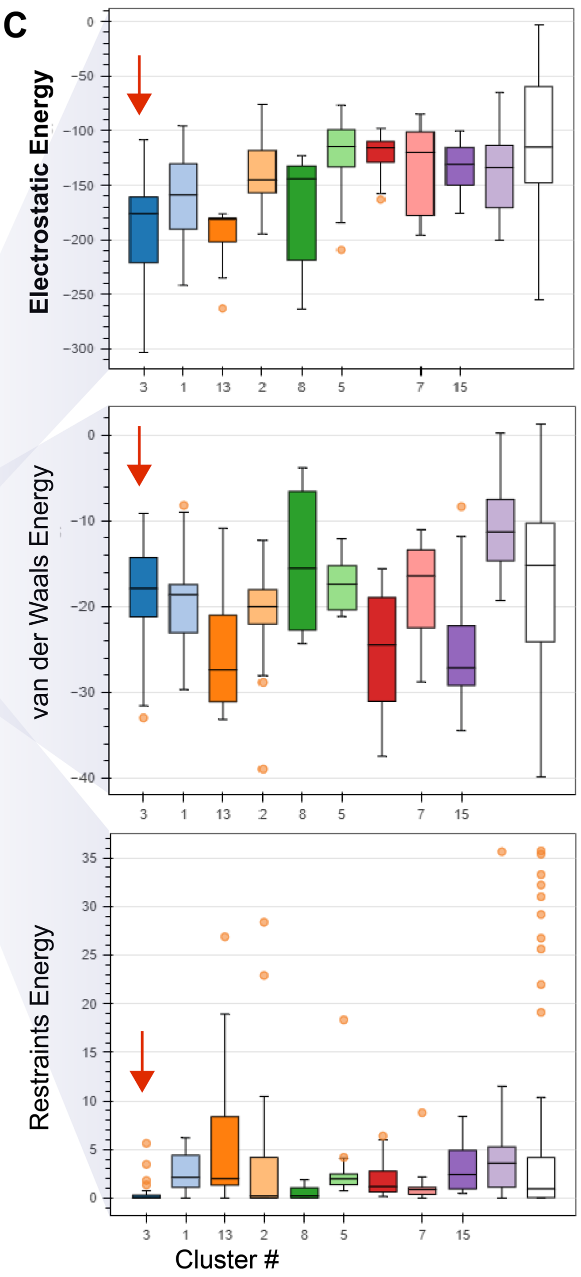

D

**Best cluster (Cluster #3)**

| | |
|---|---|
| HADDOCK score | -59.7 +/- 5.3 |
| Cluster size | 17 |
| RMSD from the overall lowest-energy structure | 1.9 +/- 0.4 |
| Van der Waals energy | -21.6 +/- 6.9 |
| Electrostatic energy | -243.8 +/- 41.2 |
| Desolvation energy | 10.7 +/- 3.0 |
| Restraints violation energy | 0.0 +/- 0.1 |
| Buried Surface Area | 1104.8 +/- 94.3 |
| Z-Score | -1.5 |

#### Supporting figure S1 [Related to Figure 2]: The workflow for the creation and analysis of the homology model of the ELMO1•SifA co-complex.

**A**. HADDOCK (High Ambiguity Driven protein-protein DOCKing) [HADDOCK2.4 web server HADDOCK Web Server (uu.nl)] is used for analyzing ELMO1•SifA interaction, which revealed several possible structures of the co-complex.
**B**. HADDOCK scoring is performed according to the weighted sum HADDOCK score of various parameters (HADDOCK2.2 scoring function; see Methods).
**C-D**. Key properties [Electrostatic, top; van der Waals, middle; and restraints energy, bottom] of the top 10 clusters are shown as whisker plots. Red → marks the properties of the best cluster (based on parameters shown in D). Cluster 3 models were chosen for further analysis based on the Z-score (-1.5; see panel D), which indicates how many standard deviations from the average this cluster is in terms of score (the more negative the better). Results indicate strong electrostatic energy as a major parameter contributing to the selection of the cluster for the formation and/or stabilization of the co-complex.

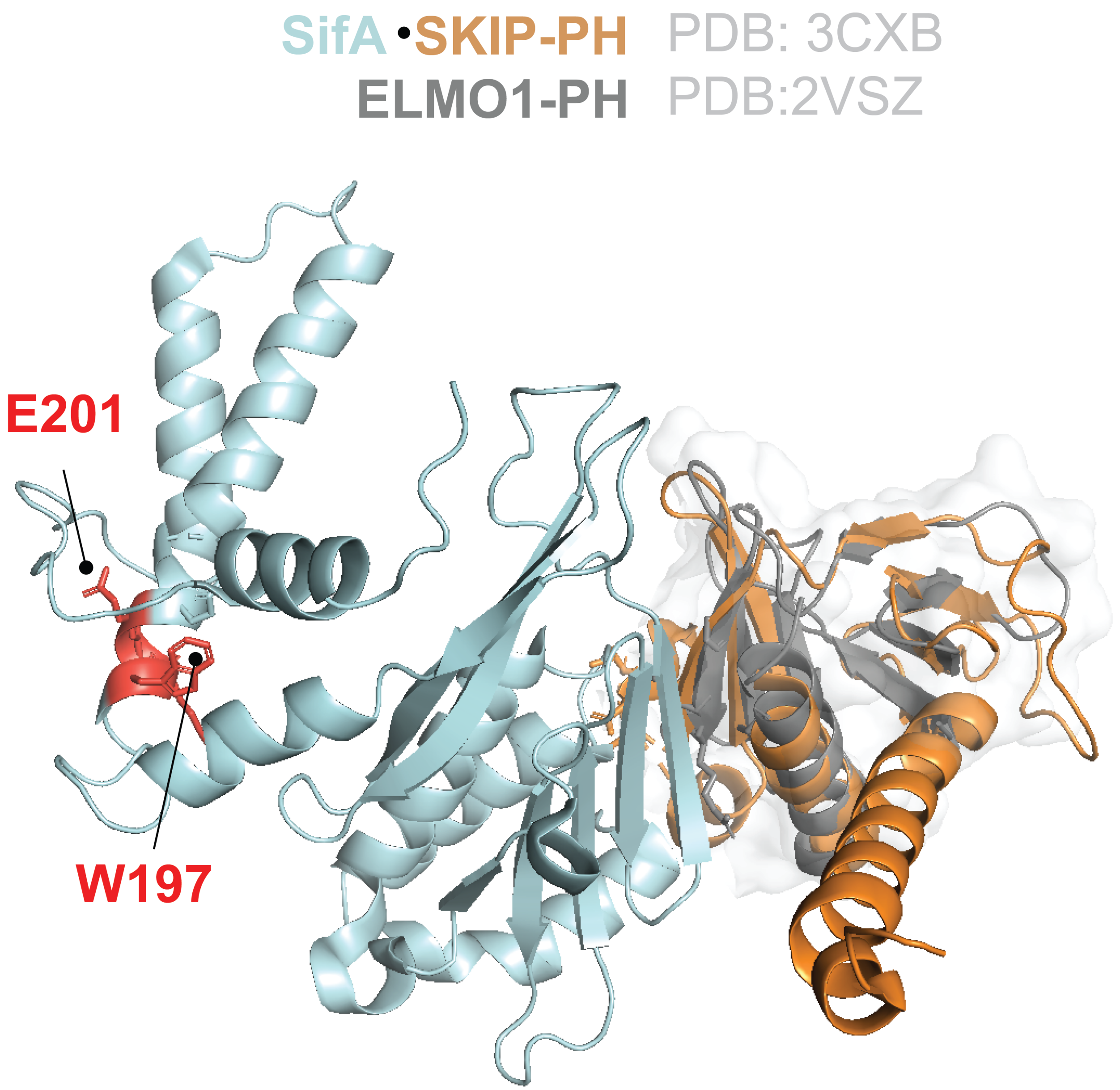

**Supporting figure S2 [Related to Figure 2]: W197 and E201 in *W*xxx*E* motif in SifA is not predicted to participate in the ELMO1•SifA interaction.**

Homology model of the ELMO1(gray)•SifA(turquoise) complex, generated by superimposing the solved structure of the ELMO1-PH (PDB:2VSZ) on that of the SKIP-PH (golden) in complex with SifA (PDB:3CXB). The executive root mean square deviation (RMSD) of the model is 1.528. The tryptophan (W) and glutamate (E) within the WxxxE-motif in SifA are highlighted in red.

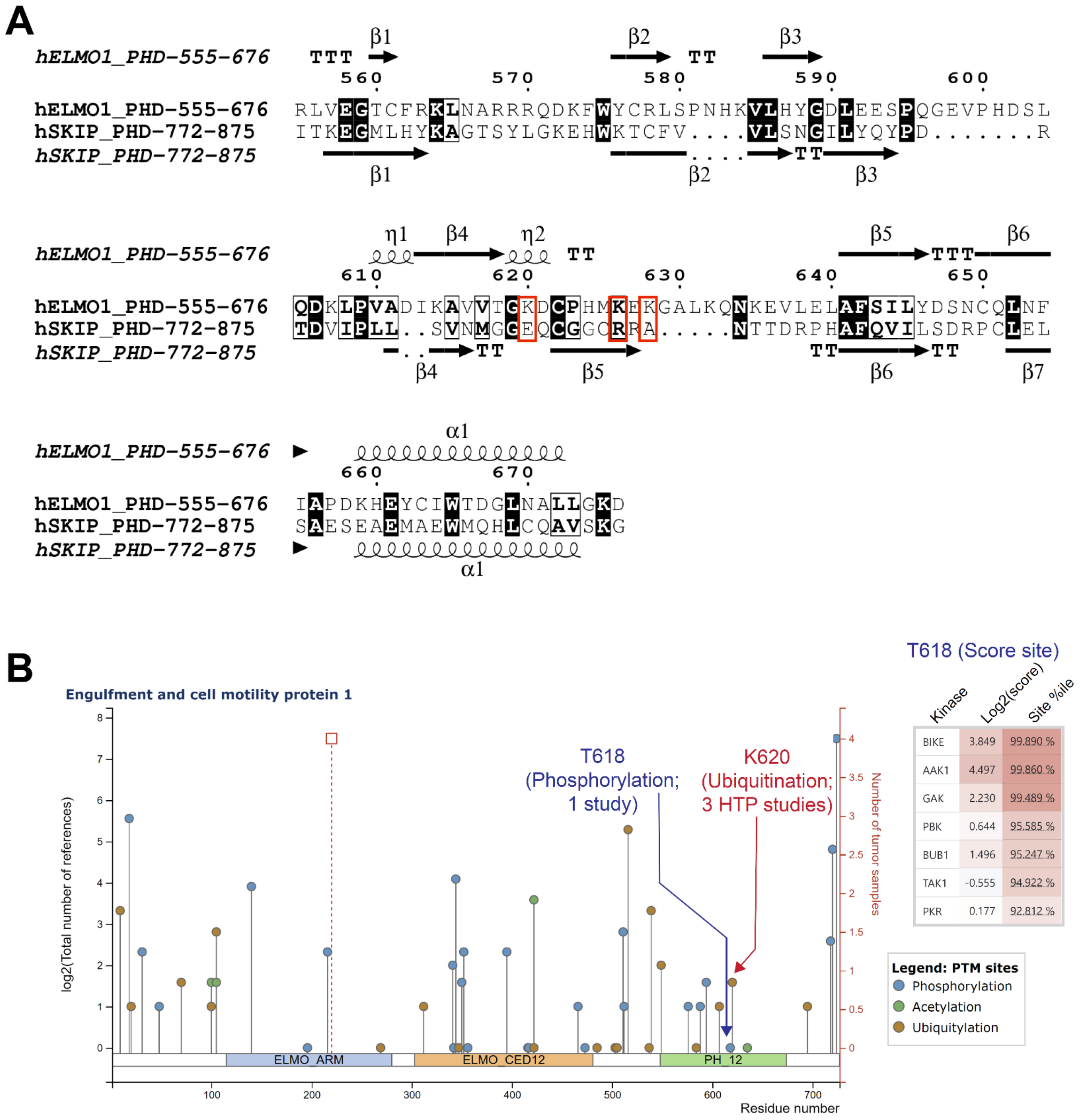

**Supporting figure S3 [Related to Figure 2]: A polar lysine triad on ELMO1 binds SifA.**

**A:** An alignment of the sequence corresponding to the PH domains of ELMO1, and SKIP is shown, along with secondary structures. Conserved residues are shaded in black, similar residues in gray box. Three lysine residues on ELMO1 were identified as critical for binding with SifA are marked with red box and were mutated in this work to confirm that they are essential for the ELMO1-SifA interaction.

**B**. Post-translational modifications identified at or near the lysine triad is shown, as curated at phosphosite.org (https://www.phosphosite.org/homeAction). K620 is ubiquitinated, whereas T618 is phosphorylated. No disease-causing SNPs or somatic mutations were found to directly impact the lysine triad.

```
MULTIPLE SEQUENCE ALIGNMENT

Human_ELMO1_AF398885.1        541------------QPEILELIKQQRLNRLVEGTCFRKLNARRRQDKFWYCRLSPNHKV
Chimp_ELMO1_XM_009452903.3    541--RPILELKEKIQPEILELIKQQRLNRLVEGTCFRKLNARRRQDKFWYCRLSPNHKV
Cow_ELMO1_NM_001113227.2      541------------QPEILELIKQQRLNRLVEGTCFRKLNCRRRQDKFWYCRLSPNHKV
Horse_ELMO1_XM_023639524.1    541--RPILELKEKIQPEILELIKQQRLNRLVEGTCFRKLNCRRRQDKFWYCRLSPNHKV
Pig_ELMO1_XM_021079240.1      541------------QPEILELIKQQRLNRLVEGTCFRKLNCRRRQDKFWYCRLSPNHKV
Mouse_ELMO1_AF398883.1        541------------QPEILELIKQQRLNRLVEGTCFRKLNARRRQDKFWYCRLSPNHKV
Rat_ELMO1_NM_001395578.1      541------------QPEILELIKQQRLNRLVEGTCFRKLNARRRQDKFWYCRLSPNHKV
Z-fish_ELMO1_XM_009294309.3   541--RPILELREKIQPEIMELIKQQRLNRLCEGTCFRKISSRRRQDKFWYCRLSPNHKV
Zenopus_ELMO1_XM_018095011.2  541------------QPEIIELIKQQRLNRLVEGTCFRKLNSRRRQDKFWYCRLSPNHKV

Human_ELMO2_AF398886.1        541-------------------IKQQRLNRLCEGSSFRKIGNRRRQERFWYCRLALNHKV
Human_ELMO3_NM_024712.5       541-------------------------LRLCEGTLFRKISSRRRQDKLWFCCLSPNHKL
                                                     ** **: ***:. ****:::*:* *: ***:

Human_ELMO1_AF398885.1        LHYGDLEESPQGEVPHDSLQDKLPVADIKAVVTGKDCPHMKEKGALKQNKEVLELAFSIL
Chimp_ELMO1_XM_009452903.3    LHYGDLEESPQGEVPHDSLQDKLPVADIKAVVTGKDCPHMKEKGALKQNKEVLELAFSIL
Cow_ELMO1_NM_001113227.2      LHYGDLEESPQGEVPHDSLQDKLPVADIKAVVTGKDCPHMKEKGALKQNKEVLELAFSIL
Horse_ELMO1_XM_023639524.1    LHYGDLEESPQGEVPHDSLQDKLPVADIKAVVTGKDCPHMKEKGALKQNKEVLELAFSIL
Pig_ELMO1_XM_021079240.1      LHYGDLEESPQGEVPHDSLQDKLPVADIKAVVTGKDCPHMKEKGALKQNKEVLELAFSIL
Mouse_ELMO1_AF398883.1        LHYGDLEESPQGEVPHDSLQDKLPVADIKAVVTGKDCPHMKEKGALKQNKEVLELAFSIL
Rat_ELMO1_NM_001395578.1      LHYGDLEESPQGEVPHDSLQDKLPVADIKAVVTGKDCPHMKEKGALKQNKEVLELAFSIL
Z-fish_ELMO1_XM_009294309.3   LHYGDLEESPQGEVPHDSLQDKLPVADIKAVVTGKDCPHMKEKGALKQNKEVLELAFSVL
Zenopus_ELMO1_XM_018095011.2  LHYGDLEESPQGEVPHDSLQDKLPVADIKAIVTGKDCPHMKEKGALKQNKEVLELAFSIM

Human_ELMO2_AF398886.1        LHYGDLDDNPQGEVTFESLQEKIPVADIKAIVTGKDCPHMKEKSALKQNKEVLELAFSIL
Human_ELMO3_NM_024712.5       LQYGDMEEGAS-PPTLESLPEQLPVADMRALLTGKDCPHVREKGSGKQNKDLYELAFSIS
                              *:***:::. .     :** :::****::*::*******::**.: ****:: *****:

Human_ELMO1_AF398885.1        YDSN---CQLNFIAPDKHEYCIWTDGLNALLGKDMMSDLTRNDLDTLLSMEIKLRLLDLE
Chimp_ELMO1_XM_009452903.3    YDSN---CQLNFIAPDKHEYCIWTDGLNALLGKDMMSDLTRNDLDTLLSMEIKLRLLDLE
Cow_ELMO1_NM_001113227.2      YDSN---CQLNFIAPDKHEYCIWTDGLNALLGKDMMSDLTRNDLDTLLSMEIKLRLLDLE
Horse_ELMO1_XM_023639524.1    YDSN---CQLNFIAPDKHEYCIWTDGLNALLGKDMMSELTRNDLDTLLSMEIKLRLLDLE
Pig_ELMO1_XM_021079240.1      YDSN---CQLNFIAPDKHEYCIWTDGLNALLGKDMMSDLTQNDLDTLLSMEIKLRLLDLE
Mouse_ELMO1_AF398883.1        YDSN---CQLNFIAPDKHEYCIWTDGLNALLGKDMMSDLTRNDLDTLLSMEIKLRLLDLE
Rat_ELMO1_NM_001395578.1      YDSN---CQLNFIAPDKHEYCIWTDGLNALLGKDMMSDLTRNDLDTLLSMEIKLRLLDLE
Z-fish_ELMO1_XM_009294309.3   YESD---EYLNFIAPDKHEYCVWTDGLNALLGKEMSSEFTRSDMDTLLNMEMKLRLLDLE
Zenopus_ELMO1_XM_018095011.2  YDSN---CQLNFIAPDKHEYCIWTDGLNALLGKDMISDLTRNDMDTLLSMELKLRLLDLE

Human_ELMO2_AF398886.1        YDPD---ETLNFIAPNKYEYCIWIDGLSALLGKDMSSELTKSDLDTLLSMEMKLRLLDLE
Human_ELMO3_NM_024712.5       YDRGEEEAYLNFIAPSKREFYLWTDGLSALLGSPMGSEQTRLDLEQLLTMETKLRLLELE
                              *: .      ******.* *: :* ***.****. * *: *: *:: **.** *****:**

Human_ELMO1_AF398885.1        NIQIPDAPPPIPKEPSNYDFVYDCN----  727
Chimp_ELMO1_XM_009452903.3    NIQIPDAPPPIPKEPSNYDFVYDCN----  742
Cow_ELMO1_NM_001113227.2      NIQIPDAPPPIPKEPSNYDFVYDCN----  727
Horse_ELMO1_XM_023639524.1    NIQIPDAPPPIPKEPSNYDFVYDCN----  742
Pig_ELMO1_XM_021079240.1      NIQIPDAPPPIPKEPSNYDFVYDCN----  727
Mouse_ELMO1_AF398883.1        NIQIPDAPPPIPKEPSNYDFVYDCN----  727
Rat_ELMO1_NM_001395578.1      NIQIPDAPPPIPKEPSNYDFVYDCN----  727
Z-fish_ELMO1_XM_009294309.3   NIQIPEVPPPIPKEPSNYDFVYDCN----  740
Zenopus_ELMO1_XM_018095011.2  NIQIPDAPPPIPKEPSNYDFVYDCN----  727

Human_ELMO2_AF398886.1        NIQIPEAPPPIPKEPSSYDFVYHYG----  720
Human_ELMO3_NM_024712.5       NVPIPERPPPVPPPPTNFNFCYDCSIAEP  720
                              *: **: ***:*  *:.::* *. .
```

**Supporting figure S4 [Related to Figure 2]: The basic 'patch' in the β5-β6 loop within the PH domain, and the triad of lysines that are predicted to bind SifA, are evolutionarily conserved in ELMO1.**

A sequence alignment of the ELMO1 PH domain region and C-terminal residues from various species, as well as the human ELMO2 and ELMO3 isoforms is shown. Green boxes indicate the previously determined DOCK180-binding sites(16); red solid box within the green box indicates the canonical 'PxxP'-motif, which is the SH3 domain binding site. The β5- β6 loop region is highlighted with a red lined box. Black arrowheads highlight the three lysine (K) residues that were predicted and validated as important residues for binding SifA.

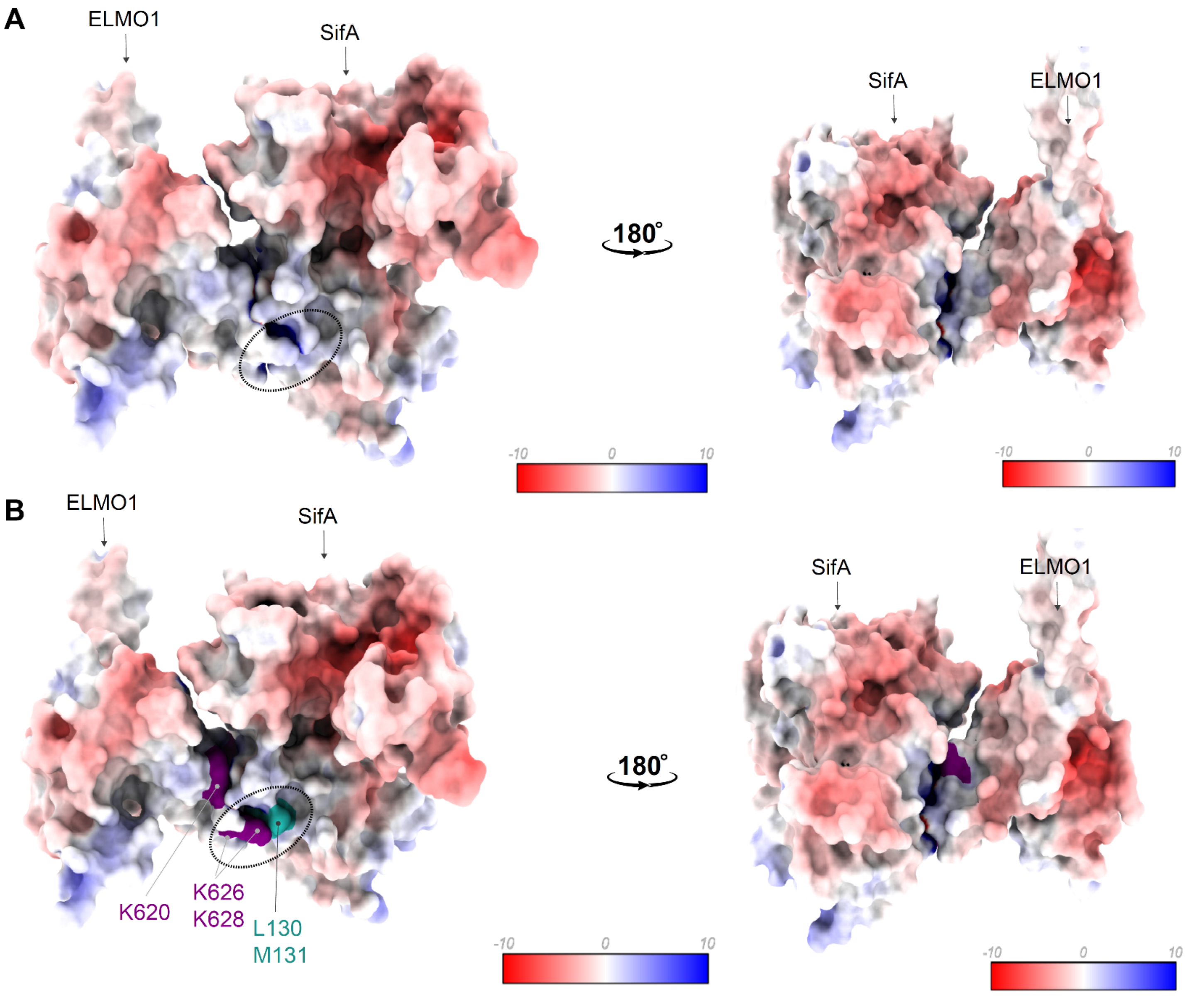

**Supporting figure S5 [Related to Figure 2]: APBS (Adaptive Poisson-Boltzmann Solver)-derived surface electrostatics of the ELMO1•SifA co-complex.**

Electrostatic surface potential (kT/e) for an all-side chain model of the ELMO1•SifA co-complex, calculated using adaptive Poisson-Boltzmann solver (APBS), an NIH-supported software (9) [https://www.poissonboltzmann.org/] for the estimation of electrostatic forces for large bimolecular assemblages. Panels display APBS output data, as visualized using Chimaera. Volume surface coloring was set in the default range of -10 (red), through 0 (white), to +10 (blue) kT/e, where negatively charged surfaces are red (-10 kT/e) and positively charged surfaces are blue (+10 kT/e). Top (unlabeled; **A**) and bottom (labeled; **B**) panels show the same co-complex, from two directions (left- and right-side views). In the most energetically favorable orientation, charged residues Lys(K)628 and Lys(K)626 on (ELMO1) bring hydrophobic residues Met(M)131 and Leu(L)130 on SifA into proximity [marked by an oval].

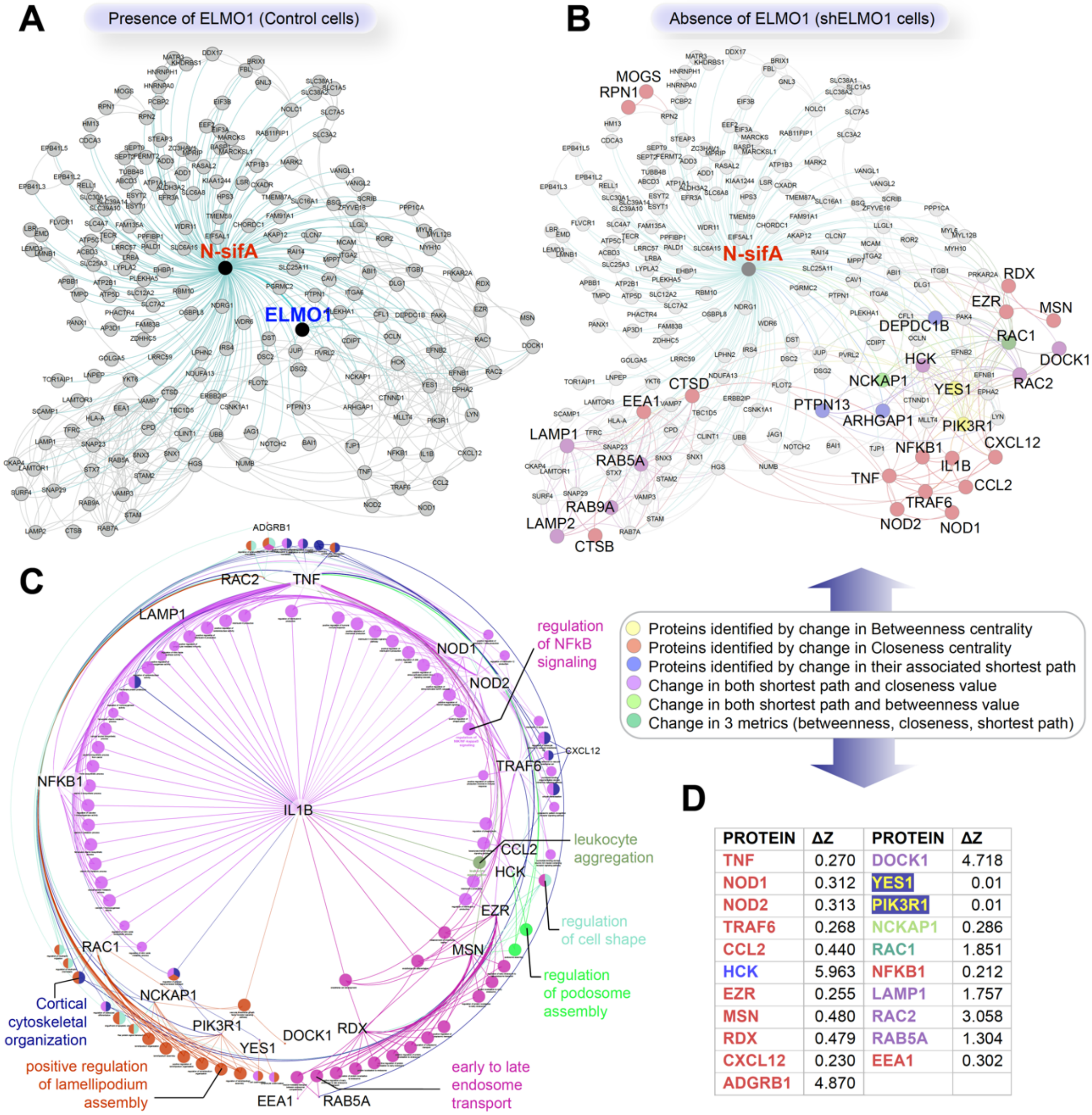

| PROTEIN | ΔZ | PROTEIN | ΔZ |
|---|---|---|---|
| TNF | 0.270 | DOCK1 | 4.718 |
| NOD1 | 0.312 | YES1 | 0.01 |
| NOD2 | 0.313 | PIK3R1 | 0.01 |
| TRAF6 | 0.268 | NCKAP1 | 0.286 |
| CCL2 | 0.440 | RAC1 | 1.851 |
| HCK | 5.963 | NFKB1 | 0.212 |
| EZR | 0.255 | LAMP1 | 1.757 |
| MSN | 0.480 | RAC2 | 3.058 |
| RDX | 0.479 | RAB5A | 1.304 |
| CXCL12 | 0.230 | EEA1 | 0.302 |
| ADGRB1 | 4.870 | | |

**Supporting figure S6 [Related to Figure 3]: Prediction of cellular consequences of depletion of ELMO1 by *in-silico* perturbation of protein-protein interaction networks.**

**A.** A protein-protein interaction (PPI) network of *SifA* and its human interactome was constructed by leveraging a previously published proximity labeling (BioID) study(2) that mapped the SifA-interactome to fetch the remaining host interactors of the SifA-interactome using the STRING database. The ELMO1●*SifA* interaction, which is the focus of current study, is highlighted. See also **Figure 3A** for workflow.

**B.** Top: A PPI network as in A, but in the absence of ELMO1 with a consequent loss of its interactions. The colored nodes highlight the topological effect of network perturbation by ELMO1 deletion, as determined by differential analysis of standard topological metrics, e.g., Betweenness, Closeness and shortest path alteration fractions. Bottom: A key with color code is presented.

**C**. Graph displays the pathway enrichment analysis of the identified proteins in B, determined using the ClueGO algorithm. Pathways with significant alterations are highlighted.

**D**. Table displays a selected list of proteins that display the greatest change in each metric, summarized as changes in Z scores. Changes > 0.1 are listed. See also **Figure 3B-C** for further analysis and interpretation.

```
SIFA_S. ENTERICA         MPITIGNGFLKSEILTNSPRNTKEAWWKVLWEKIKDFFFSTGKAKADRCLHEMLFAERAP  60
SIFB_S. ENTERICA         MPITIGRGFLKSEMFSQSAI-SQRSFFTLLWERIKDFFCDTQRSTADQYIKELCDVASPP  59
IPGB1_S. FLEXNERI        ------------------------------------------------------------  0
IPGB2_S. FLEXNERI        ------------------------------------------------------------  0
MAP_E. COLI              ------------------------------------------------------------  0
ESPM_E. COLI             ------------------------------------------------------------  0
ESPT_C. RODENTIUM        ------------------------------------------------------------  0

SIFA_S. ENTERICA         TRERLTEIFFELKELACASQRDRFQVHNPHENDATIILRIMDQNEENELLRITQNTDTFS  120
SIFB_S. ENTERICA         DAQRLFDLFCALYELSSPSCRGNFHFQHYKDAECQYTNLFIKD-GEDIPLCIVIRQDHYY  118
IPGB1_S. FLEXNERI        ------------------------------------------------------------  0
IPGB2_S. FLEXNERI        ------------------------------------------------------------  0
MAP_E. COLI              ------------------------------------------------------------  0
ESPM_E. COLI             ------------------------------------------------------------  0
ESPT_C. RODENTIUM        ------------------------------------------------------------  0

SIFA_S. ENTERICA         CEVMGNLYFLMKDRPDILKSHPQMTAMIKR-----------------RYSEIVDYPLPST  163
SIFB_S. ENTERICA         YD-------IMNRTVLCVDTQPAHLKRYSDITIKAST----------YVCEELCCLFPER  161
IPGB1_S. FLEXNERI        ----MQILNKILP---------QVEFAIPRPSFDSLSHNKLVKKILSVFNLKQRFP----  43
IPGB2_S. FLEXNERI        ---------------MLGTSFNNFGISLSH----K-RYFS------GKVDEIIRCT----  30
MAP_E. COLI              ------MFSPMT---MAGRSLVQATAQTLRPAVTRAAMQA------GTGATGMRFM----  41
ESPM_E. COLI             --------MPMN---TTGTSFSSFGISCHRENSFR-NSFL------GKNDEVIKCS----  38
ESPT_C. RODENTIUM        --------MPGT------ISSSGFGFSIAK----Q-PHSS------GQKTVIDGFF----  31

SIFA_S. ENTERICA         LCLNPAGAPILSVPLDNIEGYLYTE----LRKGHLDGWKAQEKATYLAAKIQSGIEKTTR  219
SIFB_S. ENTERICA         LLLSLSGGITFPVDLKNIKETLIAM----AEKGNLCDWKEQERKAAISSRINLGIAQADV  217
IPGB1_S. FLEXNERI        -----QKNFGCPVNINKIRDSVIDKIKDSNSGNQLFCWMSQERTSYVSSMINRSIDEMAI  98
IPGB2_S. FLEXNERI        -----MGKRIVKISSTKINTSILSSVSEQ-IGENITDWKNDEKKVYVSRVVNQCIDKFCA  84
MAP_E. COLI              -----PVQSNFVINHGKLTNQLLQAVAKQTRNGDTQQWFQQEQTTYISRTVNRTLDDYCR  96
ESPM_E. COLI             -----MGERTIHFSVRKFSGNILDTVNRQ-NTKDINGWIKDERIVYPSRVINQEIDNYCF  92
ESPT_C. RODENTIUM        -----LGTRKISFSYLRLESELMQCINLK-NEGKMNEWMREECICFVSRDVNKQLDIFAK  85
                                     .   .:   :           .   *  :*     :  ::  :

SIFA_S. ENTERICA         ILHHANISESTQQNAFLETMAMCGLKQLEIPPPHTHIPIEKMVKEVLLADKTFQAFLVTD  279
SIFB_S. ENTERICA         PPIDDAIKNKI----AAKVIENTNLKNAAFEPNYAQSSVTQIVYSCLFKNEILMNMLEES  273
IPGB1_S. FLEXNERI        -HNGVVLTSDNKKNIFAAIEKKF--PDIKLDEKSAQTSISHTALNEIASSGLRAKILKRY  155
IPGB2_S. FLEXNERI        -EHSRKIGDNLRKQIFKQVEKDY---RISLDINAAQSSINHLVSGSSYFKKKMDELCEGM  140
MAP_E. COLI              -SNNSVISKETKGHIFRAVENAL---QQPLDMNGAQSSIGHFLQSNKYFNQKVDEQCGKR  152
ESPM_E. COLI             -QKNAKISTEERQRVFSLVSHRY---QLTLDVKAAQSSINHVIMGNASFGKKIDTLCDSM  148
ESPT_C. RODENTIUM        -NNQTTIPGCVRERVFQRASFHC---GFSLDVRCAQTSTHHMILNSLYFQKKMDTLFGSA  141
                               :                      :    ::    :

SIFA_S. ENTERICA         P--STS-QSMLAEIVEAISDQVFHAIFRIDPQAIQKMAEEQLTTLHVRSEQQSGCLCCFL  336
SIFB_S. ENTERICA         S--FHG-LLCLNELTEYVALQVHNSLFSEDLSSLVETTK---NEAHH----QS-------  316
IPGB1_S. FLEXNERI        SSDMDLFNTQMKDLTNLVSSSVYDKIFNESTKVLQIEISAEVLKAVYRQSNTN-------  208
IPGB2_S. FLEXNERI        NRSVKN-DTT-SNVANLISDQFFEKNVQ-YIDLKKLR--GNMSDYITNLESPF-------  188
MAP_E. COLI              VDPITRFNTQ-TKMIEQVSQEIFERNFSG-FKVSEIK-------AITQNA----------  193
ESPM_E. COLI             SRDVKN-RTA-DCIANSLADKFYQKHIEPDIDIVKLR--NEIPDYLRCTIQA--------  196
ESPT_C. RODENTIUM        DVEVRN-QCV-RTALSSLADIFFERNVN-SIDMNKFR--DKVYDAIVQEAQRT-------  189
                                   . ::  ...  .    .
```

### Supporting figure S7 [Related to Figure 4]:

### The WxxxE-motif containing effector proteins have divergent sequences and the L130/M131 residues on SifA that bind ELMO1 are not conserved.

Multiple sequence alignment of IpgB1 [AY206439.1] and IpgB2 [AY879342.1] from Shigella flexneri, EspM [BHZP01000031.1] and Map [WP_150349845.1] from enteropathogenic E. coli, EspT [FM210026.1[ from C. rodentium, and SifA [AAA97467.1] and SifB [AF128839.1] from Salmonella enterica serovar typhimurium. The WxxxE motif (blue box) and the catalytic loop (magenta box) that was previously determined(17) as critical for the activation of small GTPases of the Rho/ family are highlighted.

### Supplementary references

1. Das, S., Sarkar, A., Choudhury, S. S., Owen, K. A., Castillo, V., Fox, S. *et al.* (2015) ELMO1 has an essential role in the internalization of Salmonella Typhimurium into enteric macrophages that impacts disease outcome Cell Mol Gastroenterol Hepatol **1**, 311-324 10.1016/j.jcmgh.2015.02.003
2. D'Costa, V. M., Coyaud, E., Boddy, K. C., Laurent, E. M. N., St-Germain, J., Li, T. *et al.* (2019) BioID screen of Salmonella type 3 secreted effectors reveals host factors involved in vacuole positioning and stability during infection Nat Microbiol **4**, 2511-2522 10.1038/s41564-019-0580-9
3. Steele-Mortimer, O., Brumell, J. H., Knodler, L. A., Méresse, S., Lopez, A., andFinlay, B. B. (2002) The invasion-associated type III secretion system of Salmonella enterica serovar Typhimurium is necessary for intracellular proliferation and vacuole biogenesis in epithelial cells Cell Microbiol **4**, 43-54 10.1046/j.1462-5822.2002.00170.x
4. Stein, M. A., Leung, K. Y., Zwick, M., Garcia-del Portillo, F., andFinlay, B. B. (1996) Identification of a Salmonella virulence gene required for formation of filamentous structures containing lysosomal membrane glycoproteins within epithelial cells Mol Microbiol **20**, 151-164 10.1111/j.1365-2958.1996.tb02497.x
5. Das, S., Sarkar, A., Choudhury, S. S., Owen, K. A., Castillo, V., Fox, S. *et al.* (2015) ELMO1 has an essential role in the internalization of Typhimurium into enteric macrophages that impacts disease outcome Cell Mol Gastroenterol Hepatol **1**, 311-324 10.1016/j.jcmgh.2015.02.003
6. den Hartog, G., Butcher, L. D., Ablack, A. L., Pace, L. A., Ablack, J. N. G., Xiong, R. *et al.* (2021) Apurinic/Apyrimidinic Endonuclease 1 Restricts the Internalization of Bacteria Into Human Intestinal Epithelial Cells Through the Inhibition of Rac1 Front Immunol **11**, 553994-553994 10.3389/fimmu.2020.553994
7. Das, S., Owen, K. A., Ly, K. T., Park, D., Black, S. G., Wilson, J. M. *et al.* (2011) Brain angiogenesis inhibitor 1 (BAI1) is a pattern recognition receptor that mediates macrophage binding and engulfment of Gram-negative bacteria Proceedings of the National Academy of Sciences of the United States of America **108**, 2136-2141 10.1073/pnas.1014775108
8. de Vries, S. J., van Dijk, M., andBonvin, A. M. (2010) The HADDOCK web server for data-driven biomolecular docking Nat Protoc **5**, 883-897 10.1038/nprot.2010.32
9. Jurrus, E., Engel, D., Star, K., Monson, K., Brandi, J., Felberg, L. E. *et al.* (2018) Improvements to the APBS biomolecular solvation software suite Protein Sci **27**, 112-128 10.1002/pro.3280
10. Pettersen, E. F., Goddard, T. D., Huang, C. C., Couch, G. S., Greenblatt, D. M., Meng, E. C. *et al.* (2004) UCSF Chimera--a visualization system for exploratory research and analysis J Comput Chem **25**, 1605-1612 10.1002/jcc.20084
11. Szklarczyk, D., Gable, A. L., Nastou, K. C., Lyon, D., Kirsch, R., Pyysalo, S. *et al.* (2021) The STRING database in 2021: customizable protein-protein networks, and functional characterization of user-uploaded gene/measurement sets Nucleic Acids Res **49**, D605-D612 10.1093/nar/gkaa1074
12. Sahoo, D., Seita, J., Bhattacharya, D., Inlay, M. A., Weissman, I. L., Plevritis, S. K. *et al.* (2010) MiDReG: a method of mining developmentally regulated genes using Boolean implications Proc Natl Acad Sci U S A **107**, 5732-5737 10.1073/pnas.0913635107
13. Qiao, L., Sinha, S., Abd El-Hafeez, A. A., Lo, I. C., Midde, K. K., Ngo, T. *et al.* (2023) A circuit for secretion-coupled cellular autonomy in multicellular eukaryotic cells Mol Syst Biol e11127 10.15252/msb.202211127
14. Sinha, S., Samaddar, S., Das Gupta, S. K., andRoy, S. (2021) Network approach to mutagenesis sheds insight on phage resistance in mycobacteria Bioinformatics 10.1093/bioinformatics/btaa1103
15. Banerjee, S. J., Sinha, S., andRoy, S. (2015) Slow poisoning and destruction of networks: edge proximity and its implications for biological and infrastructure networks Phys Rev E Stat Nonlin Soft Matter Phys **91**, 022807 10.1103/PhysRevE.91.022807
16. Komander, D., Patel, M., Laurin, M., Fradet, N., Pelletier, A., Barford, D. *et al.* (2008) An alpha-helical extension of the ELMO1 pleckstrin homology domain mediates direct interaction to DOCK180 and is critical in Rac signaling Mol Biol Cell **19**, 4837-4851 10.1091/mbc.e08-04-0345
17. Bulgin, R., Raymond, B., Garnett, J. A., Frankel, G., Crepin, V. F., Berger, C. N. *et al.* (2010) Bacterial guanine nucleotide exchange factors SopE-like and WxxxE effectors Infect Immun **78**, 1417-1425 10.1128/iai.01250-09
